# HCR-FlowFISH: A flexible CRISPR screening method to identify cis-regulatory elements and their target genes

**DOI:** 10.1101/2020.05.11.078675

**Authors:** SK Reilly, SJ Gosai, A Gutierrez, JC Ulirsch, M Kanai, D Berenzy, S Kales, GB Butler, A Gladden-Young, HK Finucane, PC Sabeti, R Tewhey

**Author notes:** denotes corresponding author. denotes contributed equally (SK Reilly, SJ Gosai).

## Abstract

CRISPR screens for cis-regulatory elements (CREs) have shown unprecedented power to endogenously characterize the non-coding genome. To characterize CREs we developed HCR-FlowFISH (Hybridization Chain Reaction Fluorescent In-Situ Hybridization coupled with Flow Cytometry), which directly quantifies native transcripts within their endogenous loci following CRISPR perturbations of regulatory elements, eliminating the need for restrictive phenotypic assays such as growth or transcript-tagging. HCR-FlowFISH accurately quantifies gene expression across a wide range of transcript levels and cell types. We also developed CASA (CRISPR Activity Screen Analysis), a hierarchical Bayesian model to identify and quantify CRE activity. Using >270,000 perturbations, we identified CREs for *GATA1, HDAC6, ERP29, LMO2, MEF2C, CD164, NMU, FEN1* and the *FADS* gene cluster. Our methods detect subtle gene expression changes and identify CREs regulating multiple genes, sometimes at different magnitudes and directions. We demonstrate the power of HCR-FlowFISH to parse genome-wide association signals by nominating causal variants and target genes.

## Introduction

Identification and functional characterization of cis-regulatory elements (CREs) – sequence modules controlling gene expression – is a critical challenge for modern genomics. While over 900,000 putative CREs have been identified via the co-occurrence of transcription factor (TF) binding, DNA accessibility, and histone modifications, few have been directly shown to regulate gene expression in their endogenous chromosomal context^1–4^. Only 7% of CREs exhibiting chromatin interactions with distal promoters interact with the nearest gene^5^. Most putative CREs remain unvalidated, with target gene(s) and quantitative effects unknown.

There is a need to better identify CREs, since over 90% of sequence variants associated with human traits and disease risk are in non-coding regions, and many are thought to impact CRE function^6^. While genetic finemapping can help pinpoint individual causal variants^7,8^, and episomal reporter assays can identify variants with regulatory potential^9–12^, methods to directly quantify the function of CREs containing variants and identify their target gene(s) are lacking. This limitation hinders our ability to identify causal variants and elucidate the mechanistic role of common variation in complex diseases and traits.

One promising approach to functionally characterizing endogenous CREs is to perturb their predicted activity using CRISPR/dCas9 to induce a targeted transcriptionally-repressive chromatin state using a tethered histone-deacetylase KRAB (CRISPRi)^13^. Generally, such approaches link single guide-RNAs (gRNAs) and the CRE’s they target in a cell to an easily observed phenotype, and are highly amenable to parallelized screening^14–17^.?

However, most CRISPR non-coding studies rely on specific cellular phenotypes downstream of gene transcription (such as cell growth or resistance to a cytotoxic drug), restricting CRE characterization to a limited subset of genes.

A more direct and generalizable strategy to assess CREs is to measure transcript abundance in response to perturbations. However, current methods to approximate transcription have suffered from significant limitations. Tagging endogenous transcripts with transgenic reporters, such as GFP, indirectly estimates transcriptional regulation via translation; furthermore, genome engineering remains a non-universally applicable and laborious task with the potential to impact downstream processes^18–20^. Single-cell RNA-sequencing of CRISPR screens directly measures transcript abundance, but are most sensitive for highly expressed genes and are currently bound in scale to relatively few gRNAs, limiting the interrogation to small portions of the genome at high costs^21^. Specialized fluorescent in-situ hybridization (FISH) techniques compatible with single-cell approaches can directly quantify transcript abundance, but current implementation in CRISPR screens has relied on proprietary, unmodifiable methods that are restricted to highly expressed genes^22^.

To address the current limitations in characterizing CREs, we developed HCR-FlowFISH, a method that combines 1) CRISPRi-mediated perturbation of CREs with 2) hybridization chain reaction (HCR), an amplified FISH method to enable quantitation of transcript abundance, for 3) flow cytometry-based single-cell measurements **(Fig. 1a)**^23^. We also developed CASA (CRISPR Activity Screen Analysis): a Bayesian inference tool for the analysis of CRISPR screens at loci with densely tiled contiguous gRNAs, which displays improved CRE identification performance over previous analysis pipelines. Similar analysis pipelines do not directly model CRE activity or their ability to both increase and decrease gene expression. Our methods address these shortcomings and are open-sourced, sensitive, and scalable to thousands of CREs and multiple genes, enabling robust screens for nearly any gene.

**Figure 1.**
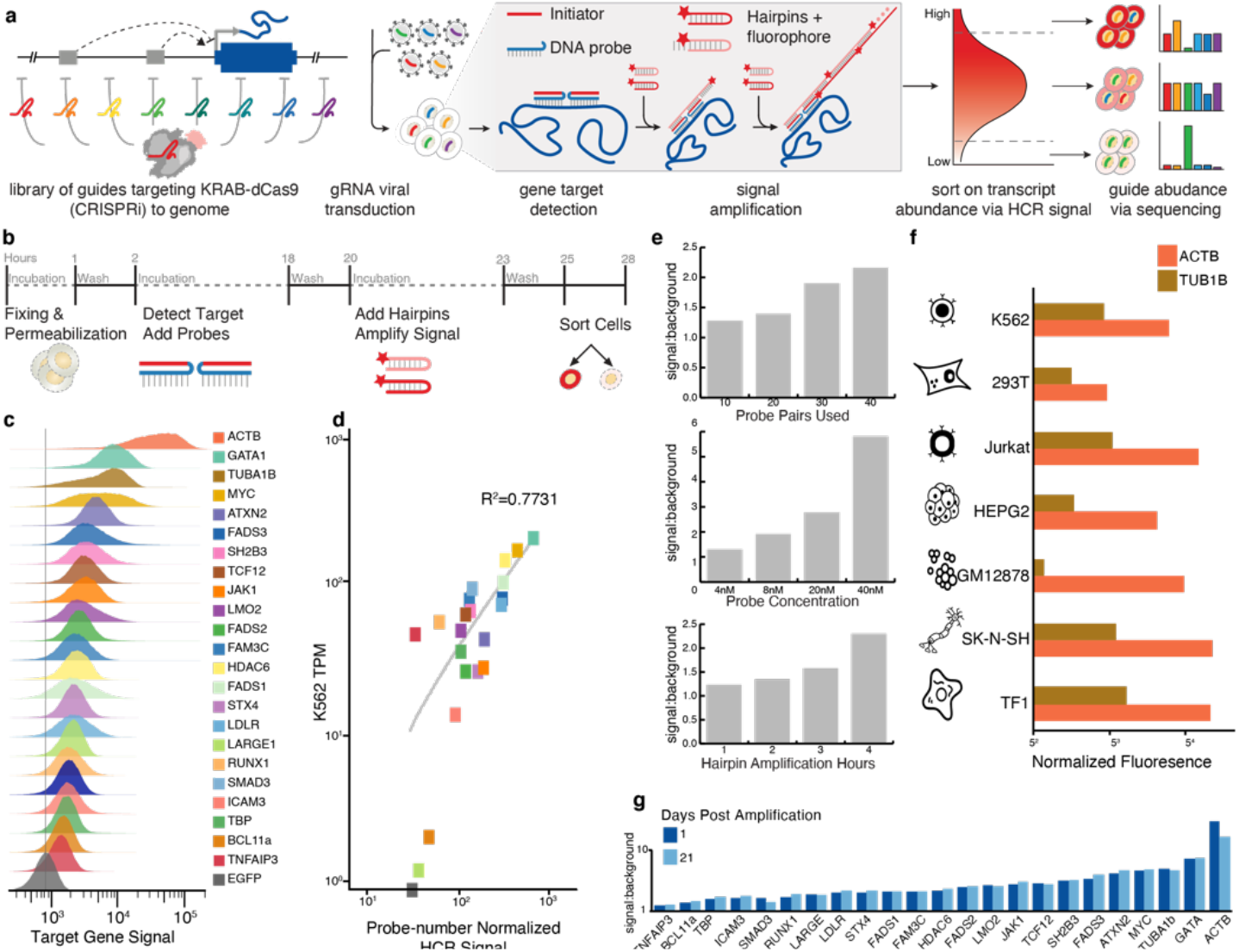
HCR-FlowFISH: a new generalizable method for transcription abundance readouts in noncoding CRISPRi screens. **(a)** Overview of HCR-FlowFISH method showing CRE identification using endogenous, CRISPRi perturbation of the genome, quantification of transcript abundance with HCR, and flow-cytometry assisted sorting to bin effector gRNAs. **(b)** Timeline of HCR-FlowFISH protocol shows shortened, 2-day protocol. **(c)** Detection of 23 transcripts and background (EGFP, non-expressed) via HCR. **(d)** Probe-number normalized HCR signal correlates with gene expression levels in K562, showcasing utility of HCR across a broad range of genes (R^2^ = 0.7731 on log10 probe-number normalized HCR signal). **(e)** Tuning of HCR signal:background ratio by increasing probe number, probe concentration, or hours of amplification increases signal to background ratio. **(f)**. Detection of *TUB1B* and *ACTB* across 6 suspension and adherent mammalian cell lines displays wide-applicability of HCR-FlowFISH. **(g)** HCR to signal to background ratio does not diminish for 21 days.

Here we apply HCR-FlowFISH to functionally characterize the regulatory landscape of ~6 Mb of the genome across 11 genes, using 275,108 gRNAs. Our endogenous perturbations revealed a more complex regulatory landscape than provided by biochemical readouts like H3K27ac and DNase I hypersensitive sites (DHSs). We detected unexpected pervasive sharing by multiple genes of regulatory elements that exhibit different magnitudes and directions of expression changes, as well as CREs regulating target genes 100kb+ away. In conjunction with massively parallel reporter assays (MPRA), we used HCR-FlowFISH to identify CREs, parse complex genetic associations, and nominate functional regulatory variants with their target gene(s).

## Results

### HCR-FlowFISH is a generally applicable, tunable, robust method for single-cell transcript quantification

We sought to establish HCR-FlowFISH as a method that enables accurate amplification of endogenous transcript levels for detection by flow cytometry. HCR quantitatively increases FISH signal by combining transcript-specific DNA probes with fluorescently labeled, metastable hairpins that form long chains, amplifying fluorescent signal once initialized by the hybridization probe. We started by modifying HCR, a method initially developed for FISH microscopy, to allow scaling to millions of single cells per experiment while providing increased transcript sensitivity and compatibility with high-throughput screening **(Fig. 1b, Methods)**^23^. We applied HCR-FlowFISH to a wide range of genes, and were able to individually detect expression of 23 transcripts relative to a negative control **(Fig. 1c)**. The assessed gene transcripts span a range of lengths (871 – 5368 bp) and expression levels (1.2–2734 transcripts per million [TPM]), with transcript levels correlating with the average probe-number normalized fluorescent signal (R^2^=0.7731) **(Fig. 1d)**.

We selected three genes to interrogate more closely, comparing HCR-FlowFISH to PrimeFlow, a proprietary RNA FISH method based on branched DNA amplification previously used for CRISPR non-coding screens^22^. For the three genes, representing low, middle, and high expression *(LARGE1 −1.5 TPM, FADS3 – 77 TPM*, and *GATA1 – 193 TPM*, respectively), we compared the signal intensity. For all three genes, we observed an improved signal-to-noise ratio (SNR) using HCR, enabling a greater dynamic range and improved detection threshold for low-abundance transcripts **(Supplementary Fig. 1a,b)**.

We investigated modifications to our HCR-FlowFISH protocol to further improve the SNR for lowly expressed genes and applicability in diverse cell types. We increased the SNR two-fold by increasing the duration of the hairpin amplification process. Increasing probe concentration and the number of probes used per target transcript increased SNR by five-fold and two-fold, respectively **(Fig. 1e)**. We successfully performed HCR for two genes across a variety of common suspension (Jurkat, GM12878, TF1) and adherent (293T, HepG2, SK-N-SH) cell lines, observing fluorescent signals concordant with cellular transcript levels, demonstrating that HCR is robust in many cell types **(Fig. 1f)**^24^. Fluorescent signals for all 23 genes tested were stable for at least 21 days **(Fig. 1g)**. Thus, signal detection and quantification can be optimized for individual gene targets and is robust across cell types.

### HCR-FlowFISH and CASA identify CREs in non-coding CRISPR screens

We next demonstrated the application of HCR-FlowFISH to measure transcript abundance as a phenotypic readout in CRISPR non-coding screens. We first assayed the *GATA1* locus which has been well studied by previous growth-based screens, allowing us to benchmark our method, while highlighting the advantages of reading out expression rather than growth^17^. Our lentiviral library contained 15,000 gRNAs targeting 90kb of sequence, comprising 278 DHSs in a 920Mb region encompassing *GATA1;* we infected into cells such that each cell received no more than one gRNA **(Fig. 1a)**.

To identify CRE-perturbing gRNAs, we then sorted doxycycline-induced CRISPRi-expressing cells and performed HCR-FlowFISH. We used different fluorescently labeled probe-hairpin combinations targeting *GATA1* and the housekeeping gene *TBP* to control for cell size and permeability, sorting for cells in the top and bottom 10% for expression **(Supplementary Fig. 2a Methods)**. DNA from the cells in each bin was isolated, gRNAs were sequenced and mapped to their genomic targets, and a guide score was calculated as the natural log ratio (low:high) expression bin gRNA abundance. Composite scores reflecting all gRNAs’ overlapping effects on a single nucleotide were also calculated **(Methods)**. Regions repressed by CRISPRi, resulting in decreased gene expression, were assigned higher composite guide scores, indicating their roles as putative CREs that enhance gene transcription.

In order to identify CREs and estimate their effect sizes from HCR-FlowFISH data, we developed CASA as a statistical framework. CASA models latent CRE activity over a disjoint sliding genomic window containing sequenced gRNAs and their propensity to alter gene expression (**Supplementary Fig. 2b**). Per replicate, we approximated posterior CRE activity scores of putative CREs and identified regions where scores diverged substantially from non-targeting control gRNAs **(Methods)**.

CASA identified three regions associated with *GATA1* expression in our pilot screen. Both composite and individual guide scores of gRNAs were strongest at the promoter of *GATA1* and two distal elements, eGATA1 and eHDAC6, previously identified in growth screens **(Fig. 2b,c) (Methods)**^17^. 1219 gRNAs were in common with a previous study **(Fig. 2a, Supplementary Fig. 3a),** and we observe strong, significant correlation both in individual gRNAs (Pearson r=0.84; p=3.4×10^-106^) and composite guide scores between studies^17^ **(Supplementary Fig. 3a,b,d)**. Correlation was much stronger in CASA-identified CREs (Spearman ρ=0.79; p=2×10^-54^) than in non-CRE regions (Spearman ρ=0.15; p=4×10^-79^) **(Supplementary Fig. 3d)**.

**Figure 2.**
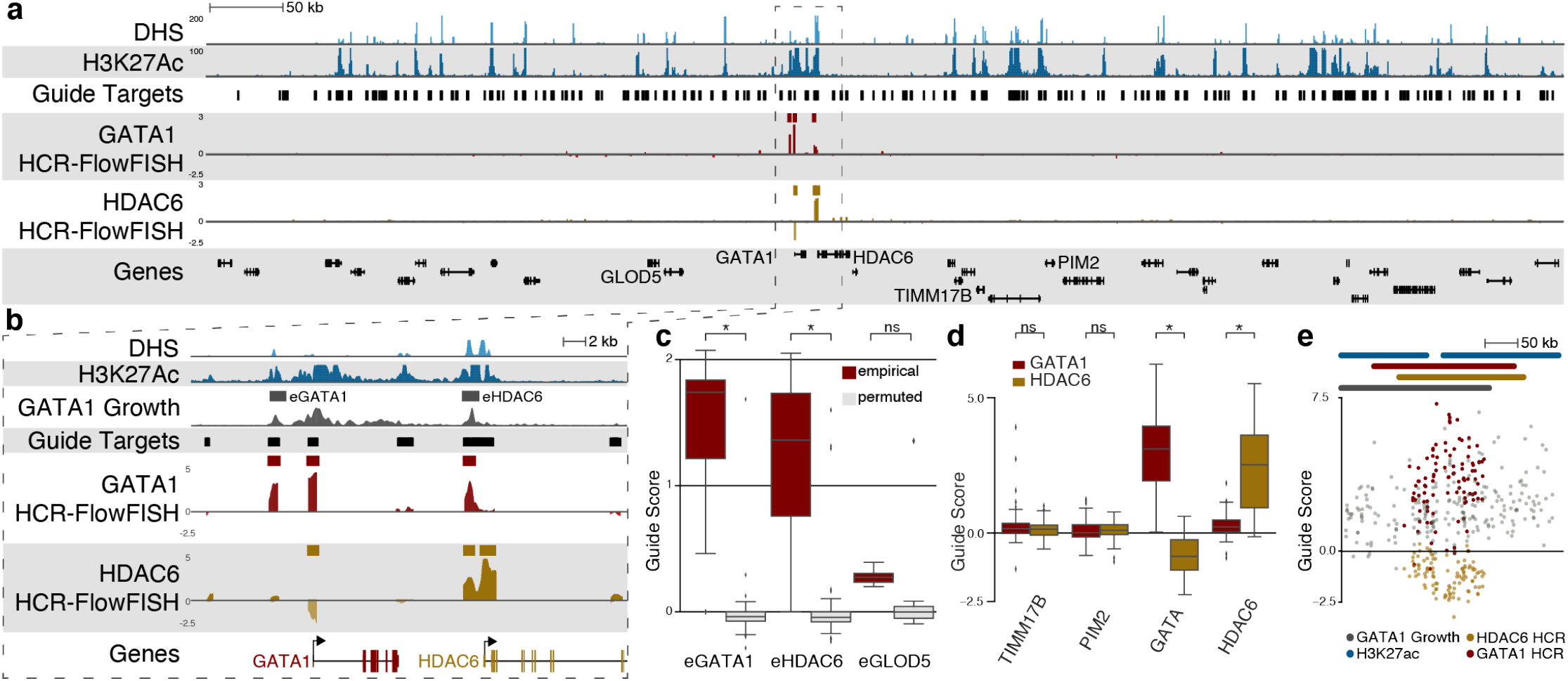
HCR-FlowFISH CRE screens on transcript abundance recapitulates growth screens at *GATA1* locus and can be extended to the *HDAC6* transcript. (**a**) HCR-FlowFISH CRISPRi Screen on a 920KB region centered on *GATA1*. Black boxes show regions targeted by guides, K562 DHS shown in light blue, H3K27ac in dark blue, composite guide scores for *GATA1* (red) and *HDAC6* (yellow) are shown. (**b**) Zoom on a 40kb region showing that CREs identified by growth screens at this locus, eGATA1 and eHDAC6, are also identified by HCR-FlowFISH, as well as the respective promoters for *GATA1* and *HDAC6.* Composite score tracks are averaged, overlapping guide scores. (**c**) HCR-FlowFISH guide scores for gRNAs in eGATA1, eHDAC6, and eGLOD5 compared to randomly permuted gRNAs. eGATA1 and eHDAC6 are identified by CASA (denoted by *, ns = not identified) by CASA. (**d**) Comparison of HCR-FlowFISH guide scores for gRNAs targeting the *GATA1* transcript (red) or *HDAC6* transcript (yellow) at promoter regions 1000bp upstream of the TSS. Guides at the *GATA1* and *HDAC6* promoter were high scoring when HCR was performed with probes for that promoter’s transcript and yielded significant CASA CREs, but not at nearby genes TIMM17B, PIM2 (* denotes p<=1×10^-5^, ns = not significant). (**e**) Individual gRNA guide scores plotted at the *GATA1* promoter locus display the opposite direction CREs for *GATA1* and *HDAC6*.

We compared HCR-FlowFISH’s performance to those of growth screens and PrimeFlow. HCR-FlowFISH identified CREs with improved specificity to growth screens: Guide scores for gRNAs within CASA-defined CREs are better separated from non-CRE-overlapping gRNAs (**Supplementary Fig. 3e, Supplementary Fig. 4a**). HCR’s improved signal to noise allows for the exclusion of spurious CREs in the *GATA1* gene body that must be removed by *post hoc* filtering in growth screens. We observed similar improvements when compared to screens using PrimeFlow (**Supplementary Fig. 4a,b**)^22^.

Because HCR-FlowFISH can be applied directly on any transcript and can interrogate more lowly expressed genes, we were able to interrogate CREs for the nearby gene *HDAC6* using the same cell library. As expected, CASA detected a CRE at the promoter of *HDAC6*, and promoter-targeting gRNAs displayed significantly increased guide scores (Mann-Whitney (MW) p < 9×10^-22^), an effect not observed at gRNAs targeting nearby promoters for *GATA1, TIMM17B,* and *PIM2* **(Fig. 2d)**.

We identified two distal CREs for *HDAC6.* Surprisingly, both overlapped CREs for *GATA1,* one overlapping the previously identified eHDAC6 element and the other overlapping the *GATA1* promoter **(Fig. 2b)**. Notably, the gRNAs perturbing the promoter of *GATA1* reduced *GATA1* expression but increased *HDAC6* expression, consistent with previous studies^17^ **(Fig. 2e)**. Our analysis at this locus highlights how HCR-FlowFISH can provide direct evidence of complex regulatory activity which eludes existing biochemical assays for chromatin structure.

### Comprehensive CRE scans of 5 loci demonstrates flexibility of HCR-FlowFISH

Next, we deployed comprehensive HCR-FlowFISH CRE-scans by selecting all high-quality gRNAs across 5 loci covering 150kb – 2.3Mb each. We used standard gRNA design software^25^ with strict cutoffs for guide specificity and efficiency to design libraries of 52,500 targeting gRNAs and 7,500 controls **(Methods)**. We first studied the moderately expressed gene *LMO2* (79.8 TPM), a regulator of hematopoiesis for which multiple CREs have been previously nominated^26,27^. CASA identifies a cluster of strong CREs 75kb upstream of *LMO2’s* promoter **(Fig. 3a)**. These CREs reside within H3K27ac peaks which have been reported to interact directly with the *LMO2* promoter and drive expression in vivo^26,27^. This stretch of CREs have been shown to recapitulate *LMO2* expression in blood cells, consistent with our use of K562 cells^28^. Additionally, we identified a canonical and non-canonical promoter, both supported by K562 CAGE^29^.

**Figure 3.**
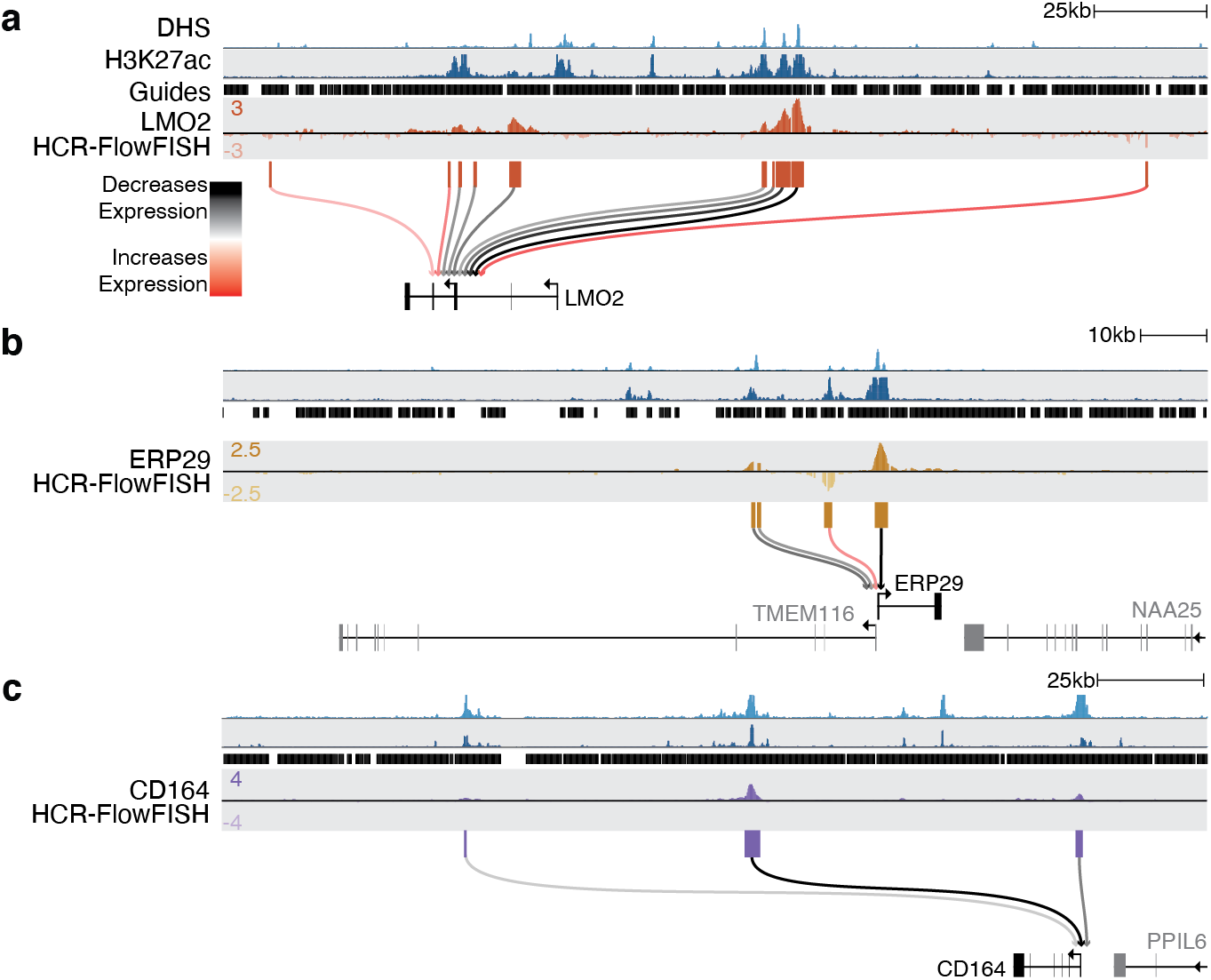
Application of HCR-FlowFISH unveils gene-specific CRE interactions at diverse loci. **(a)-(c)** Connectogram diagrams showing K562 DHS (light blue), K562 H3K27ac (dark blue), guide coverage (black), HCR-FlowFISH composite guide score tracks, and CASA CREs for LMO2 (orange), ERP29 (yellow) and CD164 (purple). CASA-derived CRE activity scores are shown as lines connecting the CRE to the target gene, and colored by effect on transcript abundance (black decreases abundance, red increases abundance). In each case, CASA identified CREs supported by K562 H3K27ac and DHS.

We also examined two other similarly expressed genes, *ERP29* and *CD164.* For *ERP29*, a reticuloplasmin, CASA identified the strongest CREs at the promoter and at distal elements within introns of the *TMEM116*gene **(Fig. 3b)**. The CREs exhibit opposing effects on *ERP29,* with the two distal CREs 19kb upstream of *ERP29* increasing transcript abundance and an 8kb upstream CRE decreasing it. This illustrates HCR-FlowFISH’s ability to detect both the strength and direction of regulatory activity. For *CD164,* a transmembrane sialomucin, we identified two CREs in addition to the promoter, −77kb and −145kb downstream of the transcription start site **(Fig. 3c)**. Notably, two closer, strong DHS/H3K27ac peaks displayed negligible effect on CD164 expression.

To test the limits of our method, we also assayed transcripts for the lowly expressed *MEF2C* (14.8 TPM) as well as *NMU* (612.8 TPM) for which small effect CREs (<10% knockdown) have been identified. *MEF2C* is a multifunctional gene important for hematopoiesis, while *NMU* acts in lymphoid cells as a signaling modulator of type 2 inflammation. For these two genes, we re-ran CASA with a more permissive activity effect threshold to call weaker-effect CREs.

As expected, CASA identified strong CREs over the *NMU* and *MEF2C* promoters **(Supplementary Fig. 5a,b).** For *NMU,* previous work has identified a CRE in a cluster of DHS sites 30.5-34kb upstream of *NMU* responsible for a 7% reduction in gene expression^21^. CASA called a CRE in the same upstream region, as well as other weak intronic CREs overlapping low levels of K562 H3K27ac **(Supplementary Fig. 5a)**. For *MEF2C,* we identified two enhancing CREs: one at its promoter, and an intronic CRE −150kb upstream. With the relaxed threshold, we identified clusters of downstream CREs ~ +40kb and +190kb and two additional intronic CREs, all supported by H3K27ac and DHS signals **(Supplementary Fig. 5b)**. These CREs range in activity scores from 40-98% of promoter repression.

Highlighting the utility of HCR-FlowFISH to aid in interpretation of non-coding genetic variation, the variant rs114694170 resides in one of *MEF2C’s* promoter-proximal CREs. We previously nominated this as a causal variant affecting platelet count by genetic fine-mapping^30^, but we lacked experimental evidence for the effect of the CRE containing this variant. Our HCR-FlowFISH maps provide direct evidence that the CRE harboring rs114694170 is a functional regulator of *MEF2C*.

### Ultradense perturbation tiling of FADS1 identifies multiple CREs and informs gRNA design

We next investigated the fatty acid dehydrogenase (*FADS*) locus (108kb surrounded *FADS1, FADS2, FADS3,* & *FEN1)* in order to comprehensively interrogate the regulatory landscape of an entire gene cluster **(Fig. 4a)**. After perturbing the *FADS* locus, sorting on *FADS1* expression, and applying CASA, we identified four clusters of CREs for *FADS1:* (i) its own promoter and a nearby but weaker 1st intron CRE; strong CREs (ii) 18kb and (iii) 25kb upstream in the introns of *FADS2* gene; and (iv) a +58kb upstream intergenic CRE between the 3’ ends of *FADS2* and *FADS3* **(Fig. 4a)**.

**Figure 4:**
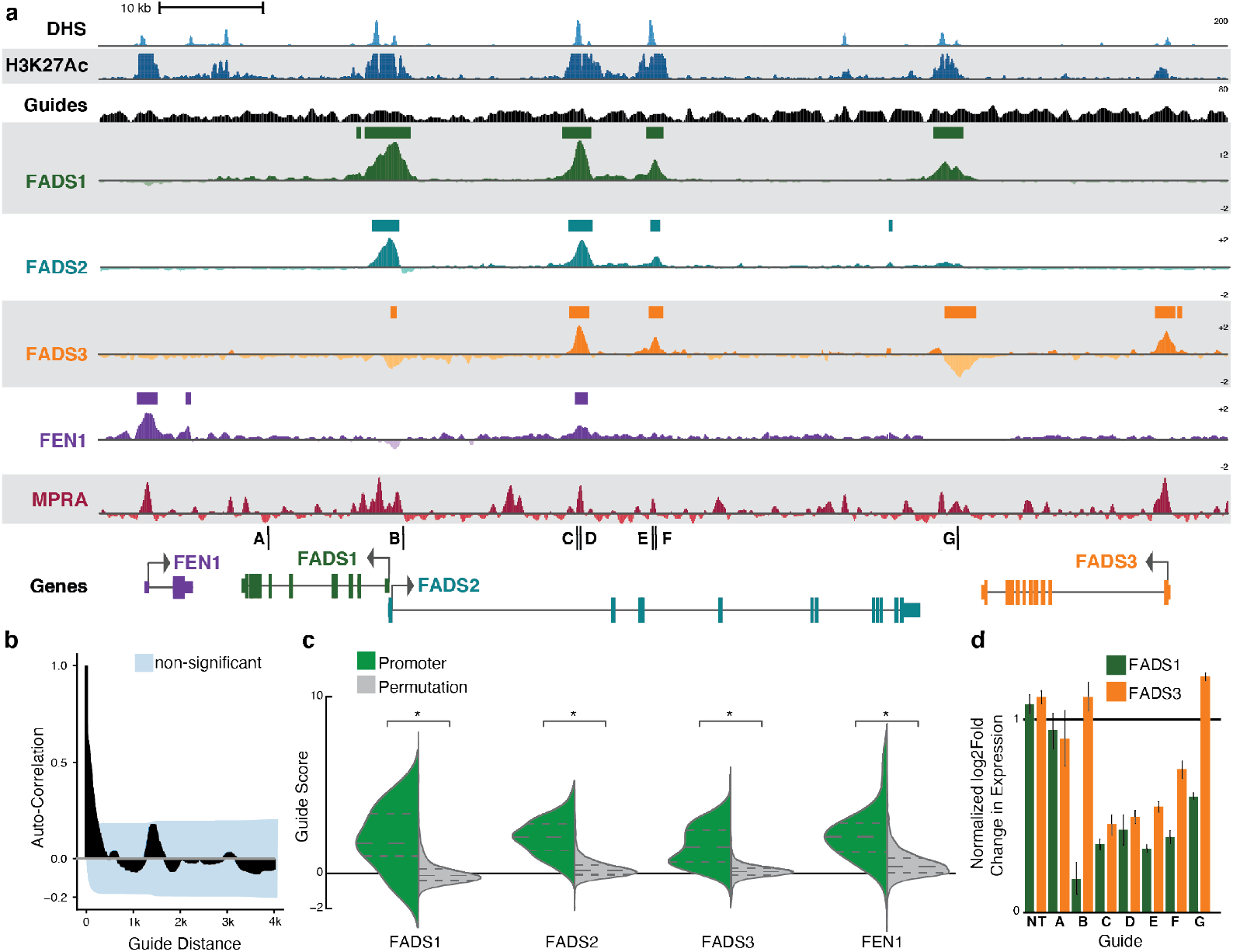
HCR-FlowFISH uncovers a complex regulatory landscape of all genes at the FADS locus. **(a)** ~100kb genomic interval surrounding the FADS locus displaying ENCODE DHS (light blue) and H3K27ac (dark blue) from K562 cells, along with the guide coverage tiled by HCR FlowFISH (black). HCR-FlowFISH composite guide scores for one replicate of FADS1, FADS2, FADS3, and FEN1 are shown. Tiling MPRA data for the same locus is included below in red. Individual gRNA binding locations for panel **(d)** are noted. **(b)** Autocorrelation plot of adjacent guides on FADS1 HCR-FlowFISH shows significant correlation of nearby guides. **(c)** Guide-wise logit scores for guides in the gene promoters shows significantly high scoring at promoters compared to an equal number of permuted guides (* = Mann-Whitney (MW) p<=1×10-10). **(d)** qPCR analysis of FADS1 and FADS3 expression changes after single-guide targetings, and a non-targeting guide (NT) corroborates transcript abundance patterns in full screen.

We also used this locus to test the relationship of gRNA design and density in a non-coding screen by densely tiling it using nearly all available gRNAs (9,554) regardless of predicted off-target quality. Even with this lenient gRNA selection criteria, we found HCR-FlowFISH experiments to be highly reproducible. For example, composite guide scores were well correlated across replicates of *FADS1* HCR-FlowFISH (Pearson r=0.98) **(Supplementary Fig. 6a)**. Furthermore, individual guide scores were highly auto-correlated between neighboring gRNAs across the locus, as expected, due to overlapping CRISPRi activity **(Fig. 4b)**.?

As expression and targeting efficiencies of gRNAs can widely vary^31^, and are shown to be associated with false positives in growth-based CRISPR screens^32^, we investigated various design metrics for association with performance measures in our screen. When comparing overlapping gRNAs between HCR-FlowFISH and *GATA1* growth screens, we observed that high representation in the cell library, rather than high gRNA specificity, appears to contribute more to the positive correlation between datasets **(Supplementary Fig. 3c).** HCR-FlowFISH results for *FADS1* supported this observation, as neither gRNA efficiency nor specificity were predictive of activity **(Supplementary Fig. 7a,b).** GC content was associated with activity, likely due to the higher GC content at promoters **(Supplementary Fig. 7c).** Dense gRNA-tiling also confirmed that modeling the strand targeted by the gRNA independently shows no differential effect, supporting a model where steric positioning of the KRAB domain is unimportant for local CRE repression **(Supplementary Fig. 7d-e)**.

We down-sampled gRNAs at the *FADS* locus to determine the minimum set required while still providing reliable CASA calls of CRE activity. The full screen has an average of 30.36 gRNAs/kb. After a 25% reduction (22.77 gRNAs/kb) noisy CREs started to be identified by CASA. These CRE calls are primarily driven by few high-scoring gRNAs in gene bodies that are outliers relative to neighboring gRNAs, creating a 1-lag autocorrelation distinctly different to that of CREs (**Supplementary Fig. 7f,g,h)**. Together, these data suggest our transcription-based assay’s less restrictive gRNA design requirements allow for additional guides, increasing accuracy of noncoding CRISPR screens.

### HCR-FlowFISH reveals the complex regulatory landscape of genes at the FADS locus

We next performed HCR-FlowFISH to quantify CRE activity for three other genes in the locus *(FADS2, FADS3* & *FEN1*), generating well-correlated replicates **(Supplementary Fig. 6a)**, and found the strongest scores at the promoter for each gene (MW p<=1×10^-10^) **(Fig. 4c).** For *FADS2,* CASA did not identify a CRE at its canonical promoter, but instead nominated an alternative promoter 12kb upstream. Only this alternative promoter was supported by CAGE data in K562 cells, suggesting it is the sole active promoter in the cell line (MW p<=1×10^-20^) **(Supplementary Fig. 6b)**.

Three non-promoter CREs identified at the *FADS* locus displayed evidence of sharing CRE activity across multiple genes **(Supplementary Fig. 8a,b)**. Perturbation of an intronic CRE 18kb downstream of the *FADS2* promoter caused a decrease in expression of all four genes tested, highlighting the extent of sharing that can occur for a single regulatory element. Other CREs were specific for only a subset of the genes at the locus. Strikingly, some regions were clearly marked by DHS and H3K27ac in K562 cells, yet perturbation of these regions did not alter gene expression at the four proximal genes measured by HCR-FlowFISH **(Fig. 4a)**, highlighting the complexity of regulatory interactions and the need for direct experimental observation. We validated the effects of several of these CREs using single gRNAs, imaging, and qPCR **(Fig. 4d, Supplementary Fig. 6c)**.

We further validated CREs identified by HCR-FlowFISH at *FADS* by orthogonal high-throughput methods. Comparing 10bp bins in HCR-defined CREs versus an equal number of randomly permuted bins not in CREs, HCR-FlowFISH identified CREs were specifically enriched for H3K27ac, DHS, PRO-Seq, GRO-seq, and CAGE data from K562 cells (MW p<=1×10^-5^ for each) **(Supplementary Fig. 9a)**. Notably, all CREs identified by HCR-FlowFISH overlapped previously identified H3K27ac peaks in K562 cells **(Fig. 4a)** and >85% of significant gRNAs directly overlapped an H3K27ac element **(Supplementary Fig. 9b)**. The composite score from HCR-FlowFISH was similar to H3K27ac signal in both position and magnitude, displaying the highest significant cross-correlation (corr=0.75) with a 0bp offset **(Supplementary Fig. 9c)**.

We found that HCR-FlowFISH was able to distinguish overlapping CREs and assign each to their target gene **(Fig. 4a,c)**. One example is an intergenic CRE for *FADS1* and *FADS3* downstream of *FADS2*, which overlaps a single H3K27ac peak and two distinct DHS peaks 1.5kb apart. While perturbation of this CRE is associated with decreases in *FADS1* expression, it also is associated with increases in *FADS3* expression.

To confirm that our screen could detect repressors as well as enhancers and promoters, and to investigate the molecular underpinnings of such elements, we targeted the *FADS3* repressive CRE identified above with a CRISPR cutting screen using 900 gRNAs overlapping the CRE. We sorted cells based on *FADS3* expression and subjected high and low expression cell populations to long-read sequencing from a single amplicon covering the CRE to identify deletions impacting gene expression **(Fig. 5a, Methods).** We applied CASA to nucleotide-resolution counts of deletions in high and low expression amplicon pools to call significant functional windows, localizing a 3000bp CRISPRi window down to a core 600bp functional region directly overlapping a DHS peak **(Fig. 5b)**. Notably, we identified three bound TFs via ChIP, along with their canonical motifs, including the transcriptional repressor REST, potentially explaining this CRE association with increased *FADS3* expression when perturbed. We also identified binding of TFs important for long-range targeting of CREs including YY1, RAD21 and CTCF **(Fig. 5c)**.

**Figure 5.**
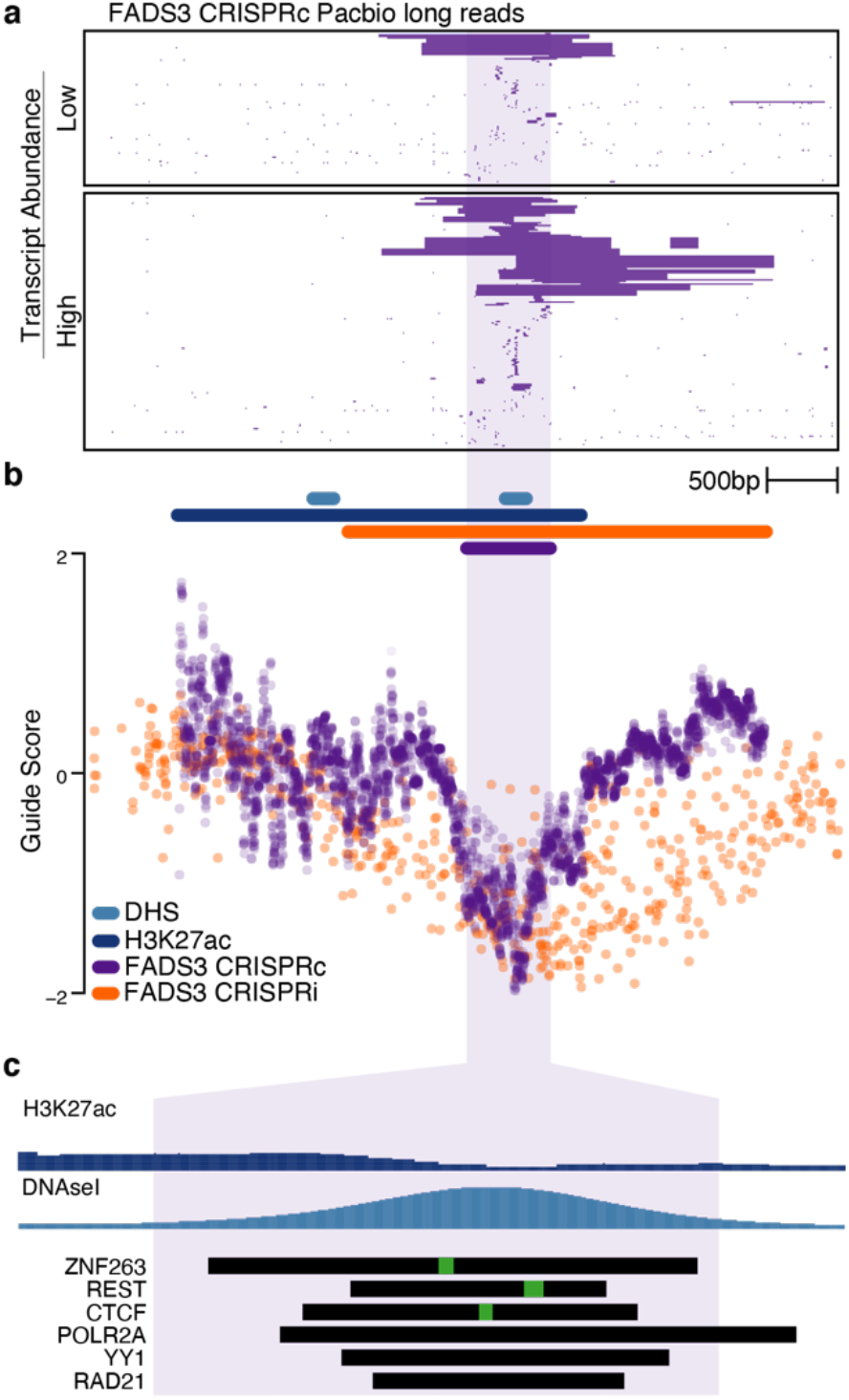
High-resolution mapping using a CRISPR cutting HCR-FlowFISH
identifies CREs at TF resolution. **(a)** Clustering of high-quality single gRNA-targeted deletion sequences (purple bars) within a 3kb CRE initially identified by the FADS3 CRISPRi screen. Deletion-bearing cells were subject to HCR-FlowFISH and sorting based on FADS3 transcript abundance. **(b)** Individual guide scores (orange) from FADS3 HCR-FlowFISH screen, overlaid with the log-odds ratio of the cumulative deletion frequencies per nucleotide in the low versus high FADS3 transcript abundance bins (purple). DHS (light blue), H3K27ac (dark blue) K562 peaks calls are shown, along with CASA CRISPR cutting CRE in purple identifying a core 500bp CRE. **(c)** Zoomed view of the CRISPR cutting CRE identifies underlying ChIP peaks for multiple transcription factors (black bars) and canonical TF motifs (inlaid green bars).

In summary, across all 7 loci that we examined with HCR-FlowFISH, our data demonstrated exceptions to several conventions of regulatory elements. We cataloged widespread regulatory element sharing between genes, something that has been described recently^19,22^. Notably, we frequently observed active distal regulators that skipped over more proximal DHS and H3K27ac regions with no functional effect. We note that neither the distance of a CRE to its target promoter nor the intensity of the DHS site were significantly correlated with CRE activity score in our dataset (**Supplementary Fig. 10a,b**). However, we did find CRE activity scores were significantly, but weakly, correlated with H3K27ac signal intensity (Spearman ρ=0.26; p=0.029) (**Supplementary Fig. 10c**).

### Combining HCR-FlowFISH and MPRA to nominate causal genetic variants and identify their effector transcripts

The FADS locus harbors a high density of independent GWAS associations for lipid metabolites and related traits, however causal SNPs and target genes have proven difficult to characterize due to a positive-selection driven elevation in linkage disequilibrium (LD) across the region^33–36^. The enzymes encoded by the genes at the *FADS* locus are necessary in biosynthesis of omega-6 and omega-3 long-chain polyunsaturated fatty acids (PUFAs), with levels of their metabolites well-reflected in blood cells^37^.

To identify functional variants that are candidate causal alleles at the *FADS* locus, we first investigated eQTLs, analyzing the 241 variants with a significant association to the expression of one or more genes in GTEx **(Fig. 6a).** Fifty-eight percent of these variants were significantly linked to multiple genes in the locus, with 57% showing the opposite direction of effect for *FADS1* and *FADS2.* With HCR-FlowFISH we determined that three eQTLs exist in CREs that regulate both genes, supporting the possibility that individual SNPs might act directly on multiple genes, rather than having multiple associations only due to tight LD to other eQTLs **(Supplementary Fig. 11a-c)**.

**Figure 6.**
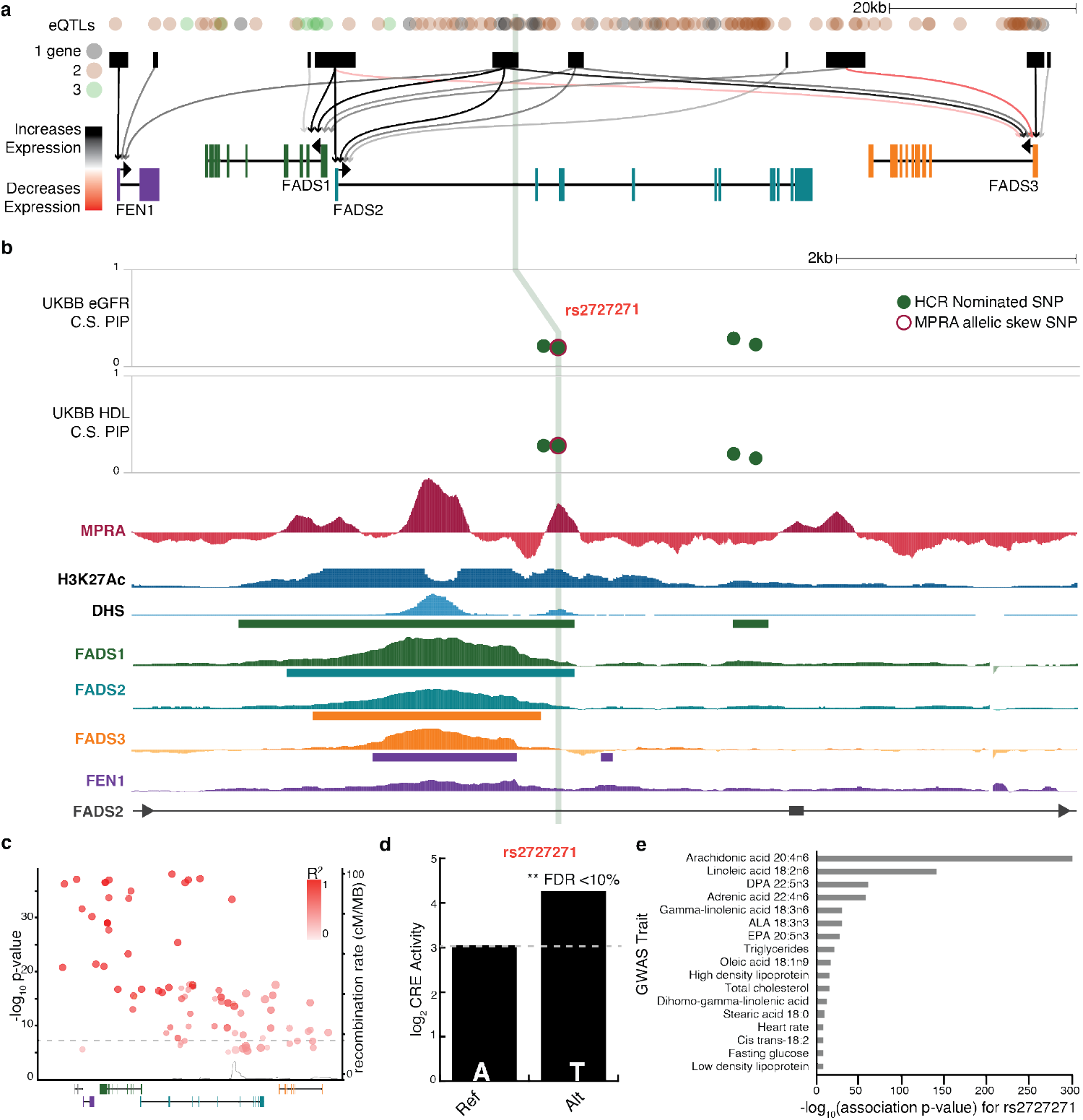
Nominating causal genetic variants and identifying their effector transcripts at the *FADS* locus. (**a**) ~110kb region surrounding the *FADS* locus showing all variants from 1000 Genomes, with notations for eQTLs and number of FADS genes they are associated with. Connectogram of all CREs identified via HCR-FlowFISH highlight complex regulatory landscape with extensive CRE sharing. (**b**) Fine-mapped credible variant sets for eGFR and HDL levels share similar posterior inclusion probabilities (PIP) due to high LD. MPRA (red), DHS (light blue) and H3K27ac (dark blue) signals, as well as composite guide scores for HCR-FlowFISH on *FADS1* (green), *FADS2* (teal), *FADS3* (orange), FEN1 (purple) at CRE within an intron of *FADS2.* Variants within a HCR-FlowFISH identified *FADS1* CRE are labeled in green, variants displaying allelic skew via MPRA are outlined in red, with location the only variant identified with both denoted by a green bar. (**c**) Total cholesterol GWAS at the *FADS* locus yields 73 genome-wide significant variants (**d**) Results of allelic skew in the MPRA shows the alternative allele drives increased CRE activity. (**e**) Many traits significantly associated with rs2727271 are direct metabolites of the *FADS1* enzyme.

To further characterize genetic variants in the *FADS* region, we combined the CRE-gene links catalogued by HCR-FlowFISH with episomal measures of CRE activity by performing an MPRA in K562 cells. We tested 200bp oligos in a 5bp sliding window along the entire *FADS* locus. Oligos with regulatory activity measured via the MPRA overlapped DHS and H3K27ac data in the region **(Fig. 4a)**, and CREs identified via HCR-FlowFISH had significantly higher MPRA scores (MW p<=1×10^-5^) **(Supplementary Fig. 9a)**. We also designed allele-specific MPRA oligos capturing every variant (3108 polymorphisms) in the 1000 Genomes Project dataset at the *FADS* locus^38^. 119 (3.8%) of the variants in the FADS region showed significant allelic skew (FDR<=10%), 51 of which occurred in HCR-FlowFISH CREs **(Fig. 6a, Supplementary Table 6, Methods)**.

We next overlaid GWAS data for blood cell composition and metabolic traits from the NHGRI-EBI GWAS catalog^39^ and Phenoscanner^40^ with MPRA and HCR-FlowFISH results in order to nominate causal variants based on allelic skew and identify their transcriptional targets. (**Supplementary Table 6)**. One of the strongest GWAS effect sizes for PUFA levels is at rs174466 (β=-0.85, p=2.5×10^-22^), which is an eQTL for 6 genes within 120kb, including *FADS1, FADS2,* and *FADS3*^41,42^. Genetic analysis alone is unable to resolve the association and which genes are affected given that 16 additional variants are in tight LD (>0.8 R^2^) with the sentinel variant **(Supplementary Fig. 12a,b)**. By deploying HCR-FlowFISH and MPRA, we showed that rs174466 is the only variant that conveys both allelic skew by MPRA and validates as a CRE by HCR-FlowFISH. The allele lies in the promoter region with regulatory potential solely acting on *FADS3*, suggesting that eQTL associations to other genes are due to LD with additional functional SNPs or trans effects. The alternate allele establishes a canonical motif for SP2, a transcriptional activator at GC box promoters, which is also bound to this location in-vivo suggesting the alternative allele for rs174466 may act to increase *FADS3* expression **(Supplementary Fig. 12a,d)**. The MPRA recapitulates this, with the alternate variant increasing activity by 0.8 log_2_ fold, notably only when the element is tested in the same orientation as its endogenous promoter **(Supplementary Fig. 12c, Supplementary Table 6)**.

Interrogating associations with total cholesterol, we again demonstrated that HCR-FlowFISH allows us to nominate individual variants from tightly-linked, genetically indistinguishable alleles, while also providing empirical evidence of the affected transcript. LD imposed similar limitations here, as 73 variants reach genome-wide significance for association (p<= 5×10^-8^) (**Fig. 6c**)^43^. Fine-mapping of cholesterol-related phenotypes, HDL levels and estimated glomerular filtration rate in the UK Biobank, nominates multiple credible sets in the region^44^,^45^. One includes a 95% credible set of four equally probable SNPs in perfect LD (R^2^=1): rs2727270, rs2727271, rs2524299, rs2072113 **(Fig. 6b, Supplementary Table 7)**. Only rs2727271 displays an allelic skew effect by MPRA (1.2 log_2_ fold) and overlaps an HCR-FlowFISH validated CRE for *FADS1* and *FADS2*.

## Discussion

The ability to interpret cis-regulatory grammar and functional organization of the genome will be essential for characterizing the mechanisms underpinning the majority of disease and trait-associated non-coding variants.

Indeed, several consortia, including ENCODE, FANTOM, and BLUEPRINT, are devoted to answering this question. In this study, we present HCR-FlowFISH, a broadly applicable approach to mapping complex regulatory interactions in nearly any expressed gene and cell system, allowing for a generalizable approach to study gene regulation.

Critically, HCR-FlowFISH allows more of the genome to be studied, as our analysis shows that low off-target or targeting efficiency scores do not confound expression-based screens, alleviating some concerns from growth-based screens regarding the global effects of off-target gRNAs^32^. At the *NMU* locus, for example, a 106kb region containing 4 previously characterized CREs is untargetable using the standard gRNA off-target filter of < 2 mismatches in the genome. The ability for future HCR-FlowFISH experiments to maintain high-quality results using a wider range of potential gRNAs will enable more comprehensive screens and improve interpretability.

CRE screens using HCR-FlowFISH reveal a rich regulatory landscape imperceptible by current biochemical assays, and offer novel insights into the regulome. We detect widespread CRE sharing across genes and surprisingly observe that CRISPRi perturbations of some CREs can impart both positive and negative effects on nearby genes. While nearly all of the CREs identified by CASA have predictive regulatory marks such as DHS and H3K27ac, classical assumptions — such as that CREs impact the nearest gene or that epigenetic signal strength correlates with gene expression — are not universally true in our endogenous screens.

Our HCR-FlowFISH screens quantify both activity and genic targets of CREs, but in its current iteration it lacks the resolution to decipher functional effects of single nucleotides. To overcome this we combined HCR-FlowFISH with MPRA to enable a powerful new paradigm for parsing genetic disease associations and nominate causal variants. However, HCR-FlowFISH is compatible with a host of different CRISPR effectors, allowing the perturbation of the genome at base pair resolution.

While this study presents detailed locus-centric, single gene-target CRE screens, HCR-FlowFISH is fully applicable to targeted genome-wide analyses. In addition, the ability to multiplex HCR can enable complex screens for multiple gene targets simultaneously. Thus, HCR-FlowFISH provides a uniquely flexible platform to innovate new technologies for the study of cis-regulatory function to facilitate the translation of genetic associations into improved mechanistic comprehension of human disease.

## Acknowledgements

C. Frieije, D. Kotliar, A. Lin, C. Myrvold, H. Metsky, B. Petros, J. Ray, C. Robinett, S. Schaffner, S. Weingarten, J. Xue, for editing and conversations about the manuscript. C. Otis, N. Pirete, and P. Rogers in the Broad Flow Cytometry Core for cytometry and sorting assistance. D. Stirling in the Broad Imaging Platform for custom scripting and assistance in image analysis. J. Ray and M. Bakalar in the Hacohen Lab for sorting and microscopy assistance. C. Fulco, J Engreitz, and E. Lander for discussion on PrimeFlow and CRISPR screens.

This work and SKR, SG, AG, SK, DB, RT were supported as an ENCODE Functional Characterization Center (UM1HG009435), a Broad SPARC grant, and the Howard Hughes Medical Institute. SR was partially supported by K99HG010669 and F32HG00922. RT is supported by R00HG008179. SG was partially supported by 4T32GM007226-41.

## Author Contributions

SKR, SJG, and RT designed experiments. SKR, AG, GBB, AGY, DB, SK, and RT performed experiments. SKR, SJG, and RT designed and performed data analysis. MK, JU, and HF performed fine-mapping analyses. SKR, SJG, AG, HF, PCS and RT contributed to the writing of the manuscript and interpretation of data.

