## Supplementary Figures for "HCR-FlowFISH: A flexible CRISPR screening method to identify cis-regulatory elements and their target genes"

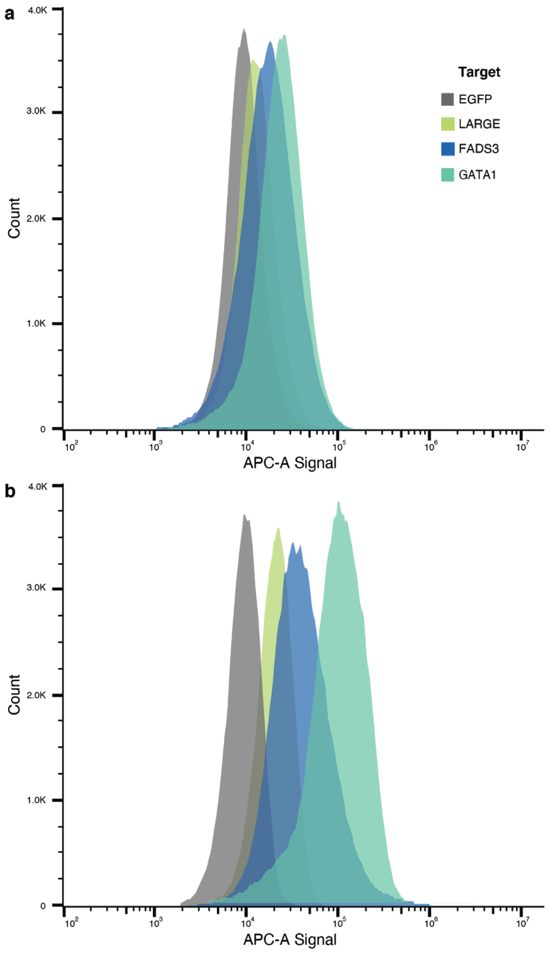


**Supplemental Fig. 1 Comparison of transcript abundance detection between HCR-FlowFISH and PrimeFlow.**
(**a**) Standard PrimeFlow protocol on three genes versus a non-expressed control eGFP (TPMs for EGFP, LARGE1, FADS3, and GATA1 are 0, 1.34, 77.70, 193.13 respectively). (**b**) Same as (**a**) but with the optimized HCR-FlowFISH protocol shows increases in signal for each transcript, but not the non-expressed negative control eGFP.

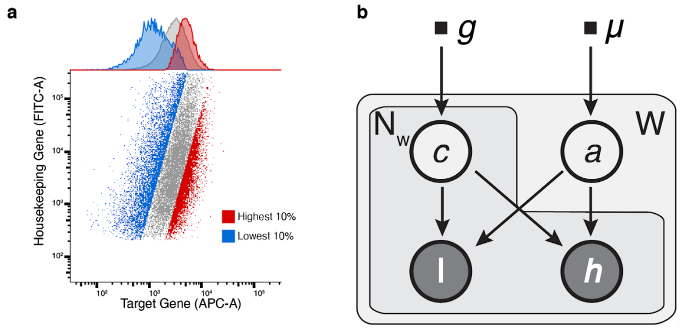


$$g \coloneqq \{\alpha, \beta\}$$

$$\mu \coloneqq \{m, s\}$$

$$c_{wn} \sim Gamma\,(\alpha, \beta)$$

$$a_w \sim \mathcal{N}\,(m, s)$$

$$l_{wn} \mid c_{wn}, a_w \sim Poisson\left(\frac{c_{wn} \times e^{a_w}}{1 + e^{a_w}}\right)$$

$$h_{wn} \mid c_{wn}, a_w \sim Poisson\left(\frac{c_{wn}}{1 + e^{a_w}}\right)$$

| Variable | Distribution | Description |
|---|---|---|
| W | None | Number of windows to test |
| $N_w$ | None | Number of gRNAs interacting with $w^{th}$ window |
| $g$ | None | Guide detection hyper priors |
| $\mu$ | None | Enhancer activity hyper priors |
| $c_{wn}$ | Gamma | Guide detection rate |
| $l_{wn}$ | Poisson | Low-expression bin counts |
| $h_{wn}$ | Poisson | High-expression bin counts |
| $a_w$ | Logit-normal | Enhancer activity score |

**Supplemental Fig. 2 CRISPRi induction, sorting schema, and construction of CASA (CRISPR Activity Screen Analysis) a generative model of CRE activity.**

(**a**) Example sorting strategy showing detection of a target transcript (*GATA1*) amplified with Alexa-647 conjugated hairpins, and a housekeeping transcript (*TBP*) amplified with Alexa-488 conjugated hairpins. The top and bottom 10% of 647:488 normalized ratio are differentially sorted. (**b**) The generative process underlying CASA (CRISPR Activity Screen Analysis) described as a plate model, explicit statistical parameterization, and variable definitions. Shaded and unshaded circles indicate observed and latent variables respectively. The variable W corresponds to the set of windows tested, while each $N_w$ arises from the set of gRNAs considered at the $w^{th}$ window.

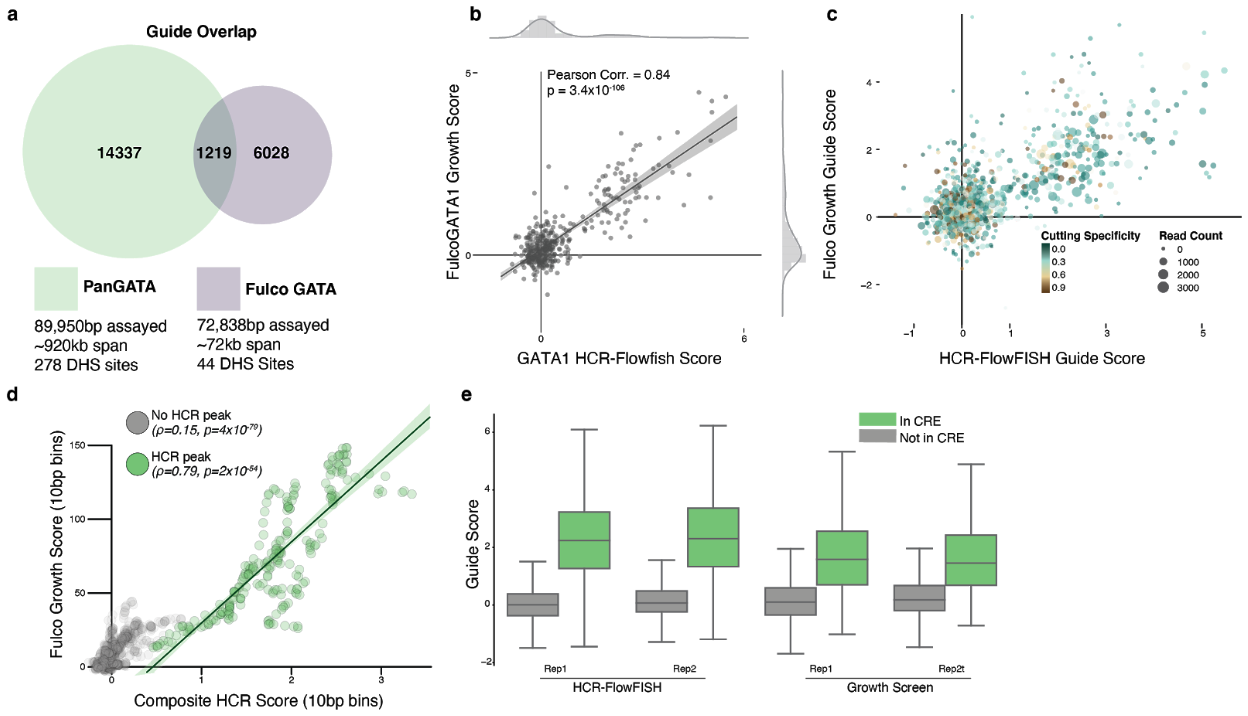


**Supplemental Fig. 3 HCR-FlowFISH screens display high similarity and increased sensitivity compared to growth screens at the *GATA1* locus.**

(**a**) Overlap of the *GATA1* guide library used in this study and Fulco et al 2016 library. (**b**) High correlation (Pearson r=0.84, $p=3.4 \times 10^{-106}$) between individual guide scores for detected gRNAs shared in the *GATA1* HCR-FlowFISH screen and the Fulco 2016 growth screen (gray shaded band is 95% confidence interval). (**c**) Guide-wise score comparison for all gRNAs shared between growth and HCR-FlowFISH screens, showing read-depth of gRNA drives correlation more than off-target effects (cutting specificity). (**d**) Comparison of HCR-FlowFISH composite guide scores compared to growth scores, both binned in 10bp regions, showing high correlation for regions in CREs identified by CASA (Spearman $\rho$ =0.79 $p=2 \times 10^{-54}$, green shaded band is 95% confidence interval). (**e**) Comparison of individual guide scores for guides shared between the HCR-FlowFISH and 2016 growth screens. The distributions scores within CREs are more distinctly separated from those without when using HCR-FlowFISH.

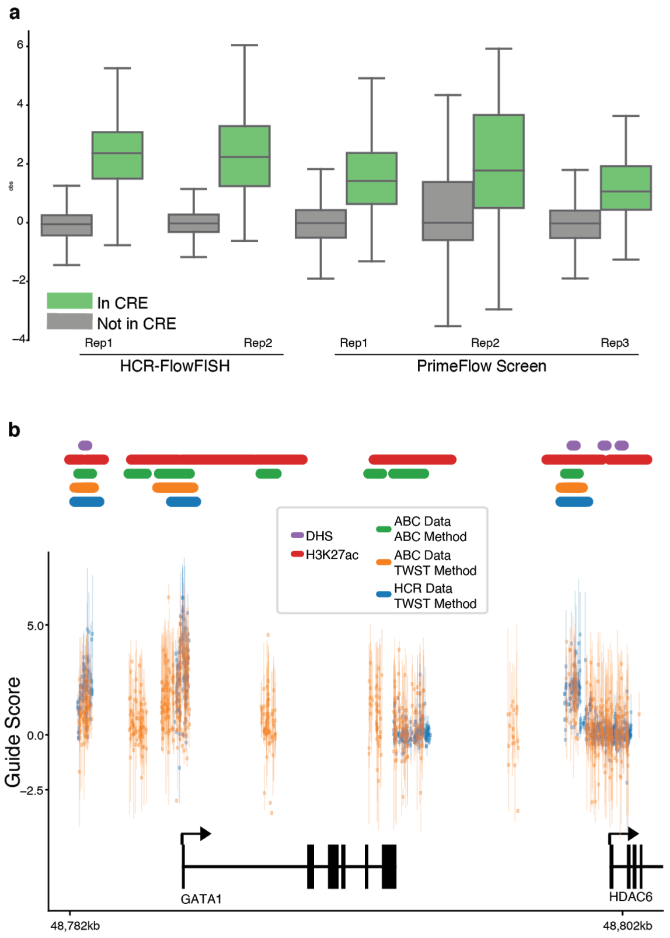


**Supplemental Fig. 4 HCR-FlowFISH and CASA enhance selectivity of CRISPRi screens at the GATA1 locus.** (**a**) HCR-FlowFISH and PrimeFlow-CRISPRi individual guide score comparison for shared guides. Guides are grouped by overlap with CASA nominated CREs. We find using HCR-FlowFISH improves separability between guide scores inside and outside of designated CREs compared to PrimeFlow. (**b**) CASA CRE identification on simplified ABC data and comparison to HCR data. CASA only considers the highest and lowest expression bins from the first PCR replicate of each CRISPRi-FlowFISH screen replicate, yet distinguishes CREs from non-specific scores induced by perturbing the GATA1 gene body, in contrast to the original analysis.

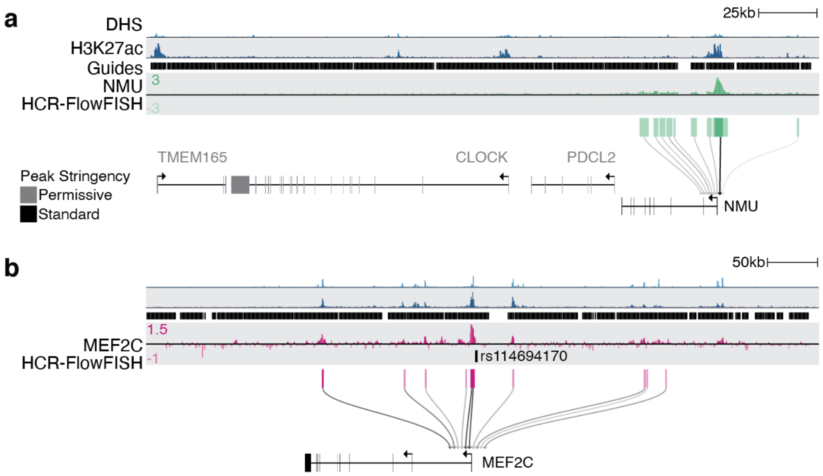


**Supplemental Fig. 5. Permissive CASA analysis identifies a CRE with modest activity on NMU and links a GWAS regulatory variant to MEF2C.**

(**a**)-(**b**) Connectogram diagrams showing K562 DHS (light blue), K562 H3K27ac (dark blue), guide coverage (black), HCR-FlowFISH composite guide score tracks, and CASA CREs calls for NMU (green) and MEF2C (pink). CASA-derived CRE activity scores are shown as lines connecting the CRE to the target gene, and colored by effect on transcript abundance (black decreases abundance, red increases abundance). For each screen, CASA odds ratios were reduced to test the limits of CRE detection, permissive CREs are displayed in lightened boxes, and were not subject to filtering at reduced odds ratio threshold **(Supplementary Table S5).**

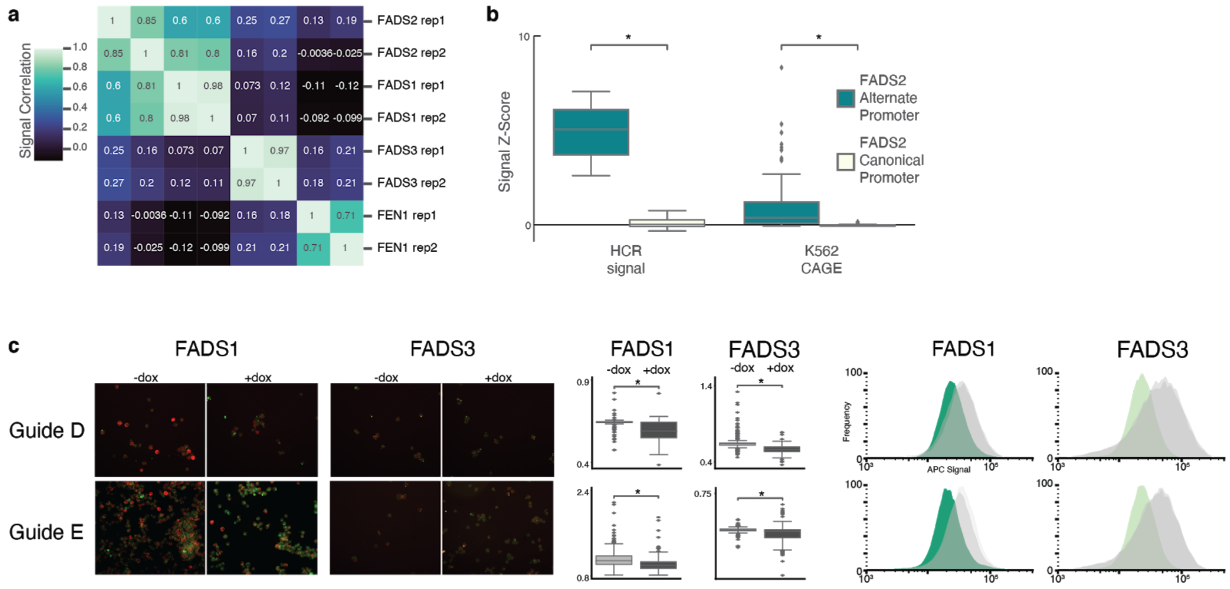


**Supplemental Fig. 6. HCR-FlowFISH produces reproducible effects in CRISPRi screens at the FADS locus.**
(**a**) Pearson correlation of replicates each of 4 HCR-FlowFISH screens for transcripts at the *FADS* locus. (**b**) HCR identifies an alternative promoter of FADS2 in K562 cells, supported by CAGE data at an alternative vs the canonical TSS (⁎ = MW $p<=1x10^{-20}$). (**c**) Imaging and cytometry on single guides from Figure 4d. Guides D and E are shown with (+, CRISPRi activated) and without (-) doxycycline. Target gene (indicated by FADS1 or FADS3 label) are labeled in Alexa-647 (red) and *TBP* labeled in Alexa-488 (green). Bar charts indicate intensity ratios of Alexa 647 to Alexa-488 staining per cell (* indicates MW $p<=1x10^{-10}$) and display reduction of the target gene when CRISPRi is activated. Histograms represent the same cell populations in images but analyzed on a flow cytometer, with doxycycline induced cells screened for for FADS1 (dark green) or FADS3 (light green) abundance compared to cells without doxycycline (grey).

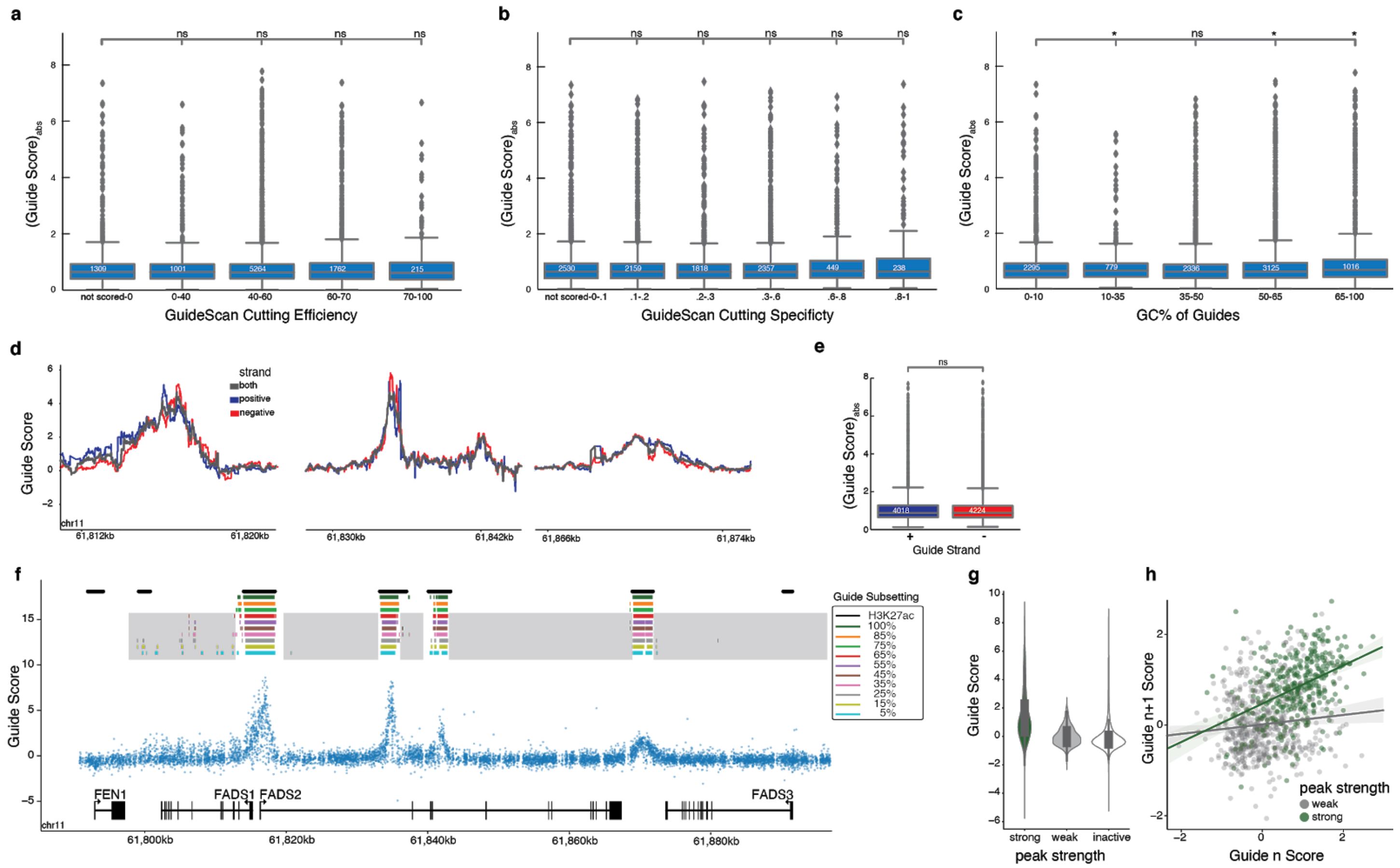


**Supplemental Fig. 7. Dense gRNA-tiling at the FADS locus informs guide design and CRE calling limitations.** (**a**)-(**c**) Absolute guide score for FADS1 HCR-FlowFISH replicates compared to Guidescan cutting efficiency or cutting specificity predictions show no correlation, but guide guide GC% displays significant correlation (ns= not significant, * = MW $p < 5x10^{-2}$). Note that for cutting efficiency and specificity guides with a 1bp mismatch to another genomic location are scored 0. (**d**) Per-nucleotide, strand-specific (blue=positive strand, red=negative strand, black=combined strands) guide score tracks calculated for FADS1 HCR-FlowFISH display similar patterns between both targeting each strand. (**e**) Absolute guide score for FADS1 HCR-FlowFISH replicates, binned by strand the guide matches to. (**f**) CASA nominated CREs based on subsets of guides using one *FADS1* HCR-FlowFISH replicate. Legend indicates the percentage of downsampled gRNAs used to identify CREs by CASA. Grey shading indicates regions where lower-confidence CRE predictions may arise after subsetting. (**g**) Gudie score distribution for guides overlapping CASA CREs with strong support when all guides are used, weak support when a subset of guides are used, or regions lacking CASA nominated CREs under any condition. (**h**) Neighboring guides in the CASA nominated CRE between FADS2 and FADS3 (green) are autocorrelated (Pearson r=0.4204, $p=2.5e^{-17}$) more strongly than guides overlapping with noisy CREs in the FADS1 gene body (grey, Pearson r=0.1041, p=0.0091).

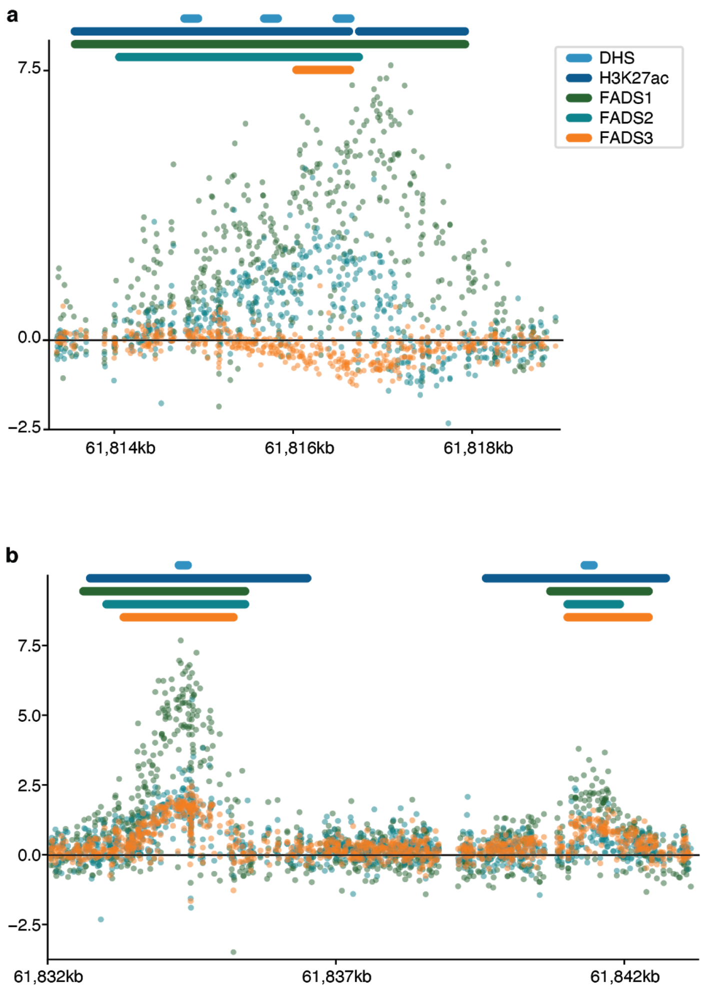


**Supplemental Fig. 8. HCR-FlowFISH and CASA reveal complex CRE sharing at the FADS locus.**
(**a**)-(**b**) Individual guide scores (points) and CASA CRE calls (bars) of HCR-FlowFISH screens for *FADS1* (green), *FADS2* (teal), *FADS3* (orange). K562 DHS (light blue) and H3K27ac (dark blue) peaks are also shown. Notably, these elements are shared between all three FADS genes. Surprisingly, perturbing the CRE in panel (**a**) results in a modest, but detectable, increase in *FADS3* transcripts, in contrast to the decreases in *FADS1* and *FADS2* transcript abundance.

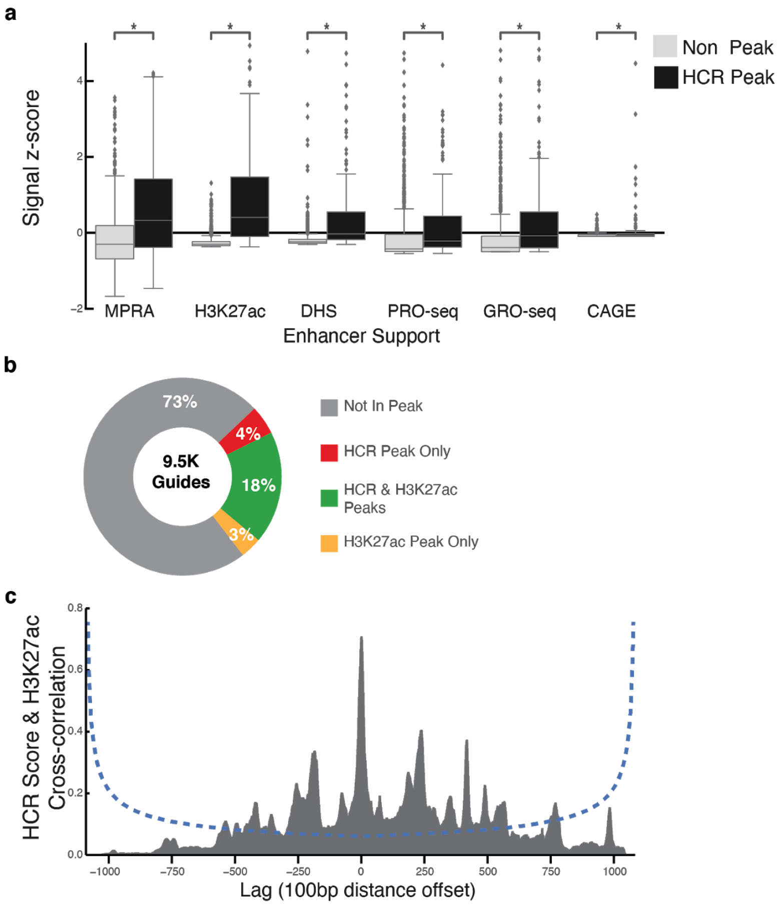


**Supplemental Fig. 9. CREs identified by CASA and HCR-FlowFISH show signatures of cis-regulatory elements.** (**a**) Six epigenetic and biochemical signatures of regulatory element show significant enrichment at HCR-FlowFISH and CASA-identified CREs at the *FADS* locus compared to non-CRE regions (* = MW $p<=1x10^{-5}$). (**b**) Percent of guides overlapping no CREs (grey), an CASA CREs region only (red), a H3K27ac peak region only (yellow), or both H3K27ac and CASA identified regions (green). (**c**) Significant crosscorrelation of H3K27ac signal and HCR-FlowFISH composite guide score (100bp binning) at the FADS locus.

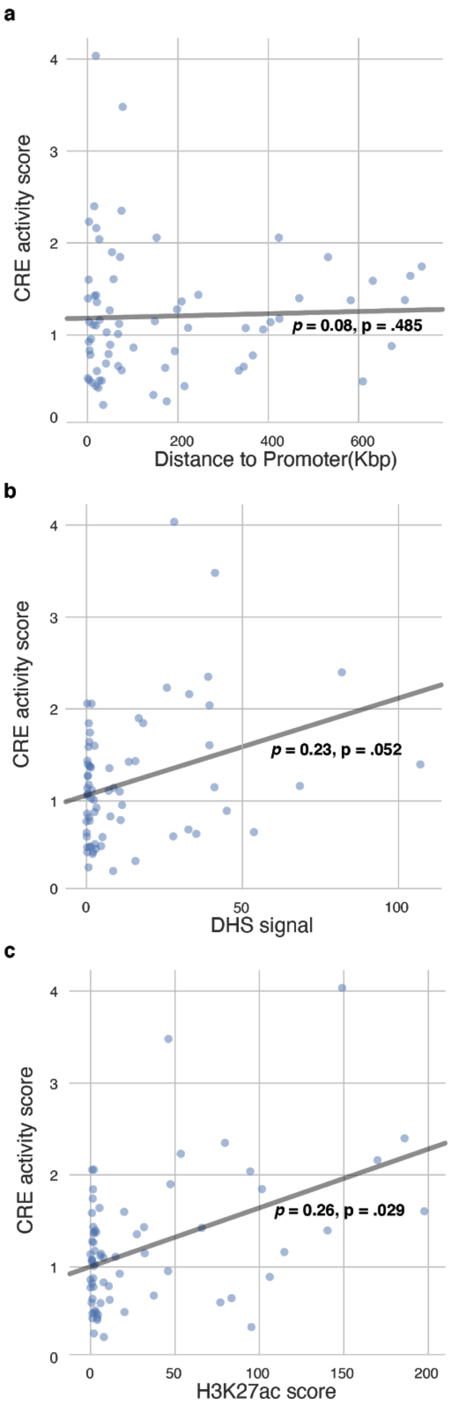


**Supplemental Fig. 10. CRE activity is weakly correlated with DHS intensity and not well correlated with distance to target gene.**

(**a**) Absolute values of CASA predicted CRE activity score (log-fold change in the sorting ratio, scaled between replicates) RE for non-promoter elements shows no significant correlation to distance from target promoter (Spearman $\rho$=0.08, p=0.485). (**b**) CRE activity scores (inclusive of promoters) compared to DHS signal intensity averaged also lacks correlation (Spearman $\rho$=0.23, p=0.052). (**c**) Weak correlation (Spearman $\rho$=0.26, p=0.029) is observed between CRE activity and H3K27ac scores.

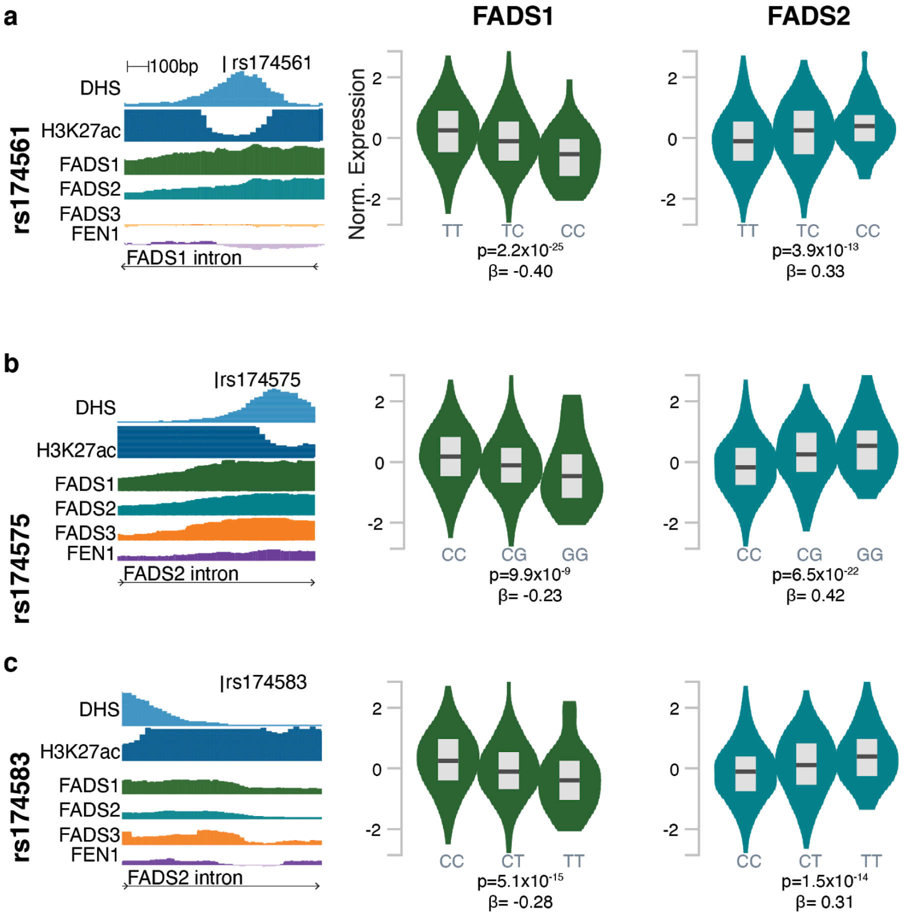


**Supplemental Fig. 11. HCR-FlowFISH identifies elements where variants have opposing regulatory effects on different genes.**

The variants: rs17451 (**a**), rs174575 (**b**), and rs174573 (**c**) overlap CREs identified by HCR-FlowFISH as regulators of both *FADS1* and *FADS2*. For each, eQTLS associated with these variants indicate significant, but opposite directions of effect for the *FADS1* and *FADS2* genes.

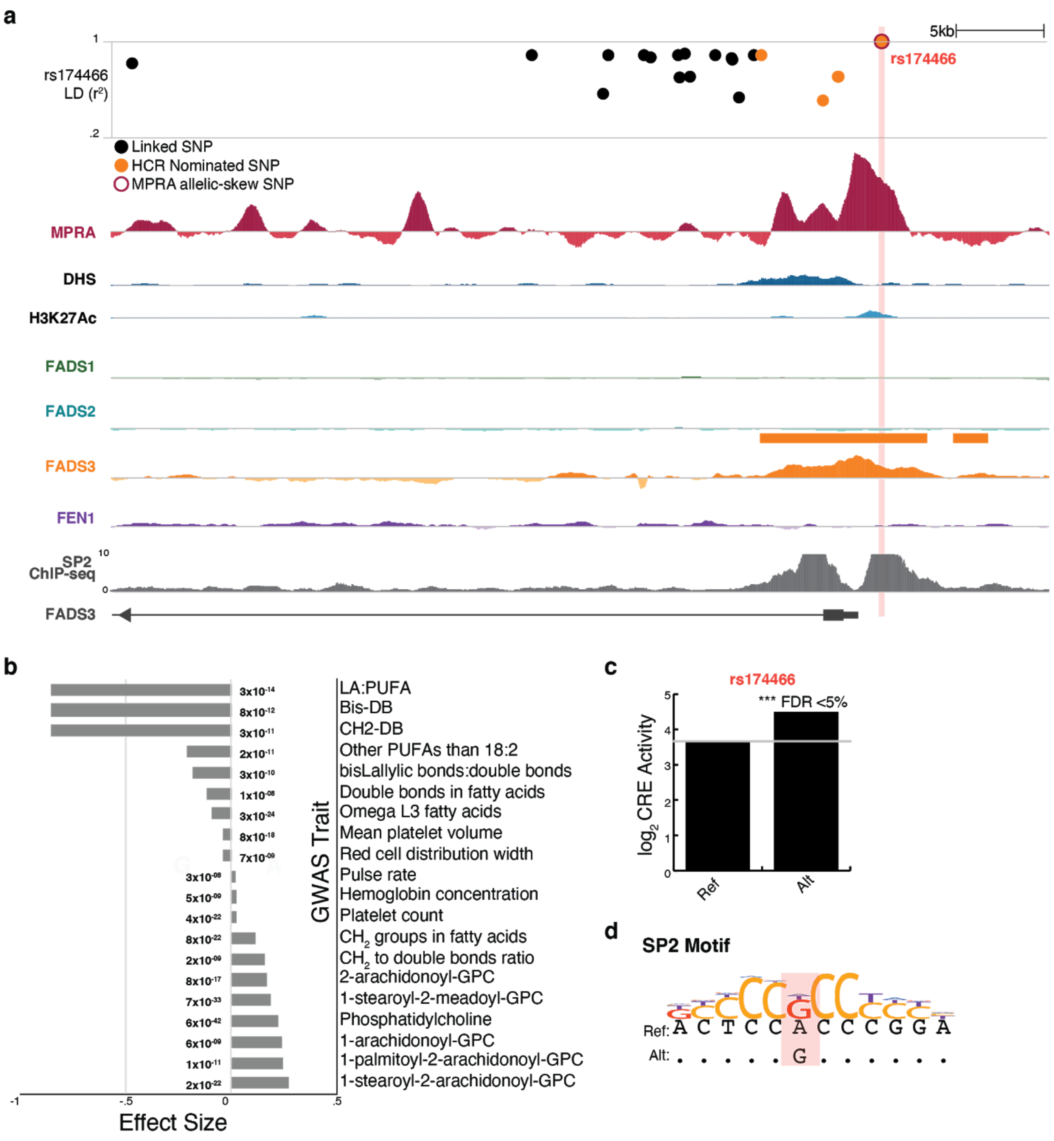


**Supplemental Fig. 12. Functional characterization nominates rs174466 as *FADS3* CRE-activity altering SNP**
(**a**) Genomic region surrounding the *FADS3* promoter, highlighting tiling MPRA signal (red) and HCR-FlowFISH composite score for *FADS3* (orange). rs174466 is denoted, along with all variants in linkage ($R^2$>=2). Variants within an HCR-FlowFISH identified *FADS3* CREs are labeled in orange, and variants displaying allelic skew from MPRA are denoted with a red outline. SP2 ChIP-Seq signal overlapping rs174466 is included in grey. (**b**) GWAS trait associations with rs174466 shows multiple overlaps with metabolic targets of FADS3. (**c**) MPRA activity for reference and alternate version of the rs174466 shows increased CRE activity on the alternate allele. (**d**) Motif for SP2 highlighting change to alternate allele better matches the canonical motif.
