## Supplementary Materials and Methods for "HCR-FlowFISH: A flexible CRISPR screening method to identify cis-regulatory elements and their target genes"

CRISPR gRNA library design, synthesis, and infection

All gRNAs and controls are reported in hg38 coordinates in (**Supplementary Table 2**). Prior to synthesis, gRNA sequences lacking a 5' guanine were appended with a G to aid in transcription efficiency. The sequence 'tatcttgtggaaaggacgaaacacc' was added before each gRNA and 'gtttaagagctatgctggaaacagcatagc' after to facilitate cloning and provide a custom Illumina read 1 primer site.

For the entire FADS gene cluster chr11:61555967-61664630 (hg19) and all DHS sites at the *GATA1* locus chrX:48,306,481-49,174,557 (hg19), gRNAS were designed using custom scripts. Briefly, all possible 20bp gRNAs with the cas9 protospacer adjacent motif "NGG" within a region surrounding were considered. gRNA efficiency scores were determined using the "Rule Set 2" method and range from 0-100, with 100 being optimal. [46] To determine the number of off-target locations, we used bowtie to map gRNAs to the human reference (hg19) with a maximum 10,000 matches, with up to three mismatches (parameters: -n 3 -l 15 -e 10000 -y --all -S) [47]. Using this set of potential mapping locations in the genome, we calculated an off-targets scores [48]. We designed 9551 and 14337 gRNAs in the *FADS* and *GATA1* gRNAs libraries respectively, with 449 and 1000 random non-targeting controls. gRNA coordinates were remapped to hg38 for reporting and plotting.

At the *CD164, ERP29, LMO2, MEF2C*, and *NMU* loci, we used the Guidescan software to design gRNAs. Guide sequences with <=1 mismatch to genome were discarded. gRNAs with specificity scores below 0.2 were discarded [25]. For each library 52,500 guides were designed to target >1Mb surrounding each gene. 1,500 non-targeting and 6,000 safe-targeting-tiling control gRNAs were included. At NMU and CD164, 1000 safe-targeting control gRNAs from Morgens et al 2017 were also included [49].

Guide oligos were synthesized by Agilent and diluted in 100ul dH20. Guide sequences were amplified using Q5 Hot Start High-Fidelity 2X Master Mix (NEB, M0494L, 10uM primers **(Supplementary Table 2)**, in a total volume of 50ul. PCR conditions were: initial denaturation 98°C – 2'; 12 cycles of 98°C – 10", 60°C – 15", 72°C – 45"), final amplification 72°C – 5"). Amplicons were purified using a 3.0x SPRI using Agencourt AMPure XP SPRI Beads (Beckman Coulter, A63881) and 2x 70% ethanol washes. SgOPTI plasmid (Addgene, 85681) was digested with BsmBI (NEB, R0580L) and subsequently purified with a 1.0x SPRI. The amplified gRNA library was cloned into SgOPTI using Gibson Assembly (NEB, E2611S) and purified with a 1.0x SPRI. The recombinant plasmid was electroporated into Endura Electro-competent cells (Lucigen, 60242-2), and grown for <16 hours at 30°C. Transformation complexity was checked by serial dilution with a 1000X minimum library size requirement. gRNA library plasmids were purified using Qiagen's Plasmid Plus MidiPrep Kit high yield protocol (Qiagen, 12945).

Lentivirus was produced as in previous studies using HEK 293T cells [50]. Transient transfections were performed with PAX2 and pCMV-VSV-G (Addgene, 35002, 8454) packaging plasmids, using the x-tremeGENE 9 DNA Transfection Agent (Sigma-Aldrich, 6365787001). Lentivirus was filtered, harvested, and frozen at -80°C until use. Cells were infected at a range of viral titres (0-200$\mu$L), to identify the amount of virus that would achieve a multiplicity of infection of 0.3. Cells were spinfected in 6M cell increments using polybrene (Sigma Aldrich TR1003G) at 1200 rpm for 45 minutes, at 37°C . Transduced cells were selected by 20ug/ml puromycin selection for 3 days. Library-scale infections performed identically to above, using the appropriate viral titre and scaling so that live cells after puromycin selection would be >1000X library size.

*Single gRNA K562 CRISPRi Cell Line Generation*
Individual gRNAs **(Supplemental Table 2)** were subjected to identical guide sequence amplification, cloning, were all performed identically to the large-scale gRNA libraries. Individual gRNA plasmids were verified by sanger sequencing.

*Cell Culture*
K562-CRISPRi cells were a gift of the Lander lab and identical to those used in previous studies [17]. Cells were grown in RPMI 1640 GluteMAX (Gibco) with 10% heat inactivated FBS (HI-FBS, Gibco). Importantly, we used a doxycycline-inducible CRISPRi system to maintain the gRNA library inactively within cells without adverse growth effects. Cells are induced with  CRISPRi induced for 24 hours with a final concentration of 1ug/ml doxycycline (VWR). Non lenti-library infected cells were periodically sorted on BFP signal when BFP/CRISPRi expression dropped below 80%. Jurkats and GM12878s were grown in RPMI 1640 GluteMAX with 15% HI-FBS, 293T cells grown in DMEM (Gibco) with 10% HI-FBS, SK-N-Sh, grown in EMEM (ATCC) with 10% HI-FBS, and TF1 cells grown in RPMI 1640 GluteMAX (Gibco) with 10% heat inactivated FBS, 2mM L-Glutamine and 2ng/ml recombinant human Granulocyte-Macrophage Colony-Stimulating Factor (GM-CSF) (Peprotech). All cells were grown at 37C and 5% $CO_2$.

*HCR-FlowFISH*
HCR probes and fluorescently labeled hairpins were purchased from Molecular Instruments using sequences listed  in **Supplementary Table 1.** Buffers were optimized for HCR-FlowFISH and prepared in lab with RNase free practices and recipes are listed below.
All samples were prepared using lo-binding plasticware (Eppendorf) where possible. All values below are listed per 5M cell aliquot. All centrifugations at 500 x G for 5 minutes unless otherwise noted.
*Cell Fixing and Permeabilization* - Centrifuge cells in conical tube, resuspend in 1ml of 4% formaldehyde in PBST (1x PBS, .1% Tween 20) and incubate at room temperature for 1 hour with rotation. Pellet cells, aspirate supernatant and wash with 1ml of PBST. Repeat for a total of 4X PBST washes. Pellet cells and resuspend cells in cold 70% EtOH and incubate at 4°C for 10 minutes, and then pellet cells.
*Expression Detection* - Wash cells 2X with .5ml of PBST, pelleting cells after each wash. Resuspend Cells in 400ml of pre-warmed probe hybridization buffer (30% formamide, 5X sodium chloride sodium citrate (SSC), 9mM citric acid pH 6.0, .1% Tween 20, 50ug/ml heparin, 1X Denhardt's solution, 10% low MW dextran sulfate). Pre-hybridize for 30 minutes at 37°C with rotation in hybridization over. During pre-hybridization, prepare 100ml of probe hybridization buffer and 2pMol and warm to 37°C. Add probe solution to sample for a final probe concentration of 4 nM, incubate overnight (15-20 hours) at 37°C with rotation.
*Excess Probe Removal* - Add 500ul of SSCT wash buffer (5X SSC, .1% Tween), and centrifuge at 750X G for 5 minutes. Remove as much supernatant as possible without disturbing the pellet, which will not be compact in first wash. Resuspend pellet with 500ul of probe wash buffer (30% formamide, 5X SSC, 9mM citric acid pH 6.0, .1% Tween 20, 50ug/ml heparin) and pellet cells. Repeat for a total of four probe washes. Resuspend cell pellet in 500ul SSCT wash buffer, incubate at room temperature for 5 min then pellet cells.
*Signal Amplification* - Resuspend cells in 150ul of pre-warmed amplification buffer (5X SSC .1% Tween, 10% low MW dextran sulfate) and pre-amplify for 30min at room temperature with rotation Prepare 15pMol each of labeled hairpins by boiling 5ml of 3mM hairpin at 95°C for 90 seconds and then allowing to cool to room temperature in the dark for >=30 minutes. Prepare hairpin mixture by adding all cooled hairpins to 100ul of

amplification buffer. Add hairpin mixture to sample to reach 60nM hairpin concentration. Incubate samples at room temperature in dark with rotation for 3 hours (*FADS1, FADS2, FADS3, GATA1, HDAC6, CD164, NMU*) to 12 hours (*ERP29, MEF2C, LMO2*). Centrifuge samples and aspirate hairpin solution. Resuspend cell pellet in 500ul of SSCT wash buffer and pellet cells. Repeat SSCT wash for a total of six washes. Resuspend cells in PBS and keep in dark at 4°C prior to sorting.

*Prime-Flow*

The PrimeFlow RNA Assay Kit was purchased from Thermo Fisher Scientific (88-18005-204) along with corresponding probes *GATA1* (VA1-20436-PF), *LARGE1* (VA1-3005734-PF), *FADS3* (VA1-6005976-PF) and eGFP (60175). Cells were stained in triplicate according to the specifications provided in the kit. The cells were analyzed on a Beckman Cytoflex identically to HCR-FlowFISH stained cells.

*Sorting and Cytometric Analysis*

Cells were filtered using a Celltrics 0.3um filter (04-004-2326, Sysmex) and diluted with PBS to an approximate concentration of 10 million cells/mL. The cells were sorted using a Sony MA900, using a 100uM chip (LE-C3210, Sony), using the 405nm, 488nm, 638nm lasers. Cells were first gated for live, single-cell, and BFP+ cells as our CRISPRi is tagged with blue-fluorescent protein.

To accurately quantify changes in *target gene* expression in CRISPRi-perturbed cells, we take advantage of the multiplexable nature of HCR by assigning different hairpin-probe combinations to different transcripts. HCR-FlowFISH uses this feature to simultaneously quantify transcripts for a target gene and the housekeeping gene *TBP* to control for differential fluorescence signal caused by cell size, permeability, and total RNA abundance in each cell. One HCR probe-hairpin-fluorophore combination is used to label *GATA1* transcripts and a second combination for *TBP*.To sort cells, we plotted FITC (*TBP* probes) vs APC (target gene probes) and gated on the highest and lowest 10% FITC/APC ratio (**Supplementary Fig. 2a)**. Each sorting bin contained a minimum of cells equal to 100x gRNA library size to maintain complexity. The collected cells were spun down at 500g for 5 minutes and frozen at -20°C until DNA isolation.

Single gRNA libraries and other non-sorted HCR-FlowFISH samples were analyzed using a Beckman CytoFLEX LX Flow Cytometer. A minimum of two replicates each of 100,000 total cells were used. We analyzed APC (target gene) and FITC (*TBP*) single on live, single-cell, BFP+ gated cells using FlowJo v9.

*DNA Isolation*

We isolated DNA from cells using the following protocol for 1M cells and scaled accordingly: Pelleted cells were thawed on ice and resuspended in 100$\mu$l of Lysis Buffer (1% SDS, 10mM EDTA, 50mM Tris-HCl, pH = 8.1), and incubated at 65°C for 3 hours. Lysed cells were cooled to room temperature with 14 units of RNase A (Qiagen, 19101) and incubated for 30 minutes at 37°C. Protein digestion was achieved by adding 8 units of Proteinase K (NEB, P8107S) before a final set of incubations, 37°C for 2 hours, followed by 20 minutes at 95°C for inactivation. DNA was purified with a 1.0x SPRI and 5x 70% EtOH washes with bead resuspension in each individual ethanol wash. Final elution in 80ul of DNase-free water from the beads was preceded by a 65°C incubation for 5 minutes.

*Library preparation and sequencing*

Guide sequences were amplified directly into sequencing libraries from all recovered genomic DNA. A maximum of 600ng genomic DNA was used per PCR reaction: 25ul of Q5 Hot Start High-Fidelity 2X Master Mix, 2.5ul of 10uM indexed forward/reverse primers **(Supplementary Table 2)**, 1ul of 10mM spermine (Sigma-Aldrich,

85590-5G), in total volume of 50ul. PCR conditions were: initial denaturation 98°C – 2'; 22 cycles of 98°C – 10", 60°C – 15", 72°C – 45"), final amplification 72°C – 5"). All reactions were pooled to maintain complexity. 600ul of amplified gRNA sequence was purified using a double sided SPRI (0.5x followed by a 1.2x SPRI) to remove primers and genomic DNA with 2x 70% ethanol washes. Libraries were quantified on a High Sensitivity D1000 ScreenTape (Agilent, 5067-5584), associated reagents (Agilent, 5067-5585), and Agilent Technologies' 2200 TapeStation. Libraries were pooled and sequenced with either Illumina MiSeq v2 50 cycle kit (15033623) or NextSeq v3 75 (15057941) cycle kit using custom read one and indexing primers **(Supplementary Table 2).**

*HCR-FlowFISH normalization and visualization*

We represent data from HCR-FlowFISH experiments in three forms: individual guide scores, composite guide scores, and CRE activity scores. The first two scores are derived directly from the data, as described in this section, while the third is inferred from a statistical model defined in the next section. Once gRNA counts were measured for each sorting bin in an experiment, we conducted a semi-random scaling procedure to equalize counts between pairs of bins. First, gRNAs which are not detected in either the low-expression or high-expression bins are discarded, Second, the larger library is scaled down by the small-to-large library ratio. Based on counts observed in the low-expression bin, $l$, and the high-expression bin, $h$, for all $N$observed gRNAs, the scaled values are:

$$l_n^{\ddagger} = l_n \frac{\min\left(\sum_{n=1}^{N} l_n, \sum_{n=1}^{N} h_n\right)}{\sum_{n=1}^{N} l_n}$$

and

$$h_n^{\ddagger} = h_n \frac{\min\left(\sum_{n=1}^{N} l_n, \sum_{n=1}^{N} h_n\right)}{\sum_{n=1}^{N} h_n}$$

Finally, we convert each continuous scaled value to a discrete count by isolating the integer parts and incrementing them up by one with probability equal to the fractional part. We use these scaled library pairs for all downstream analyses, and suppress this scaling in subsequent notation.

Next, we assign an area of effect to each gRNA as the 300 nucleotide window surrounding the predicted Cas9 cut-site for each gRNA. To visualize the data, we determine a nucleotide-wise composite guide score as the log-odds ratio (LOD-score) of the total counts observed in the low-expression bin ($l$) divided by the total in the high-expression bin for all gRNAs in the set $M_k$which affect a given nucleotide, $k$:

$$s_k = \frac{\sum^{M_k} l_n}{\sum^{M_k} h_n}$$

Therefore, positive scores indicate enhancer activity. We also report individual guide scores by calculating LOD-score for each gRNA in each experimental replicate. To aid visualization, we shifted down the composite guide scores by the median of the individual guide scores for each replicate.

*CASA: Cis-regulatory activity prediction*

We designed a hierarchical Bayesian model for CRISPR activity screen analysis (CASA) to demarcate putative CRE-gene connections using individual HCR-FlowFISH experimental replicates (Sup Fig. 2c). In our model, we assume that each gRNA has a detection rate, $c$, in a sequencing library the prior distribution:

$$P(c_n) \sim \mathrm{Gamma}(\alpha, \beta).$$

Concurrently, the latent CRE activity score of each genomic window is drawn from the prior:

$P(a_w) \sim \mathcal{N}(m, s)$.

Finally, for each genomic window, $w$, and gRNA , $n$, we assume the likelihoods of observed gRNA counts in a pair of low and high gene target expression libraries are:

$$P(l_{wn} \mid c_{wn}, a_w) \sim \text{Poisson}\left(\frac{c_{wn}e^{a_w}}{1+e^{a_w}}\right)$$

and

$$P(h_{wn} \mid c_{wn}, a_w) \sim \text{Poisson}\left(\frac{c_{wn}}{1+e^{a_w}}\right),$$

respectively. This formulation is similar to the Gamma-Poisson model of gene expression where a latent rate parameter induces an overdispersed count distribution. The key difference in our model is a logit-normally distributed CRE activity score which partitions observable gRNA detection rate between two sequencing libraries generated from the same HCR-FlowFISH experiment.

CASA determines the posterior latent CRE activity, $a$, for each genomic window, $w$, of predefined size in a locus conditioned on HCR-FlowFISH count data for the $N_w$gRNAs with overlapping areas of effect:

$$P(a_w|\mathbf{l}_w, \mathbf{h}_w)$$

Additionally, we compute $a_w$conditioned on a large set of non-targeting gRNAs to standardize latent activity measurements:

$$P(a_w - a_{NT}|\mathbf{l}, \mathbf{h})$$

and assume all $a_w$ are conditionally independent to make inference tractable.

Finally, we use the region of practical equivalence (ROPE) decision rule to determine which genomic windows compose putative CRE-gene connections [51]. Briefly, we set a symmetric ROPE centered on zero for each tested gene and calculate the 95% highest density interval (HDI) for each standardized CRE-activity prediction. Putative active bins are deemed significant when the entire 95% HDI falls completely above or below the ROPE. The absolute values of the ROPE boundaries are: $log\ 2.0$for *GATA1*, *HDAC6*, and *FADS1*; $log\ 1.6$ for *FADS2*, *FADS3*, and *FEN1*; $log\ 1.8$ for *ERP29*, and *LMO2*, *CD164*, *NMU*, and *MEF2C*. We also used low-stringency ROPE boundaries of absolute value $log\ 1.2$ and $log\ 1.4$ for *NMU* and *MEF2C*, respectively. We discard solitary bins with ostensibly significant CRE-activity which are not contiguous with at least one more significant bin. This filters out artifacts due to poor coverage or edge effects from binning. Finally, where applicable, we only visualize CRE-gene pairs with consistent support between experimental replicates.

We implemented CASA in PyMC3 [52] and use the No-U-Turn Sampling (NUTS) algorithm [53] to numerically compute posteriors. We have provided CASA in several forms to the end user, including the source code on GitHub, a public Docker environment, and a Python widget that can automatically run the analysis on the Google Cloud Platform using local data. Code to visualize results is also hosted on GitHub.

*Cis-regulatory element identification*

We report CREs based on CASA activity prediction with consistent support between experimental replicates. First, within each replicate, we merge contiguous runs of CASA nominated bins with significant CRE activity scores. Next we merge putative CREs across experimental replicates for a given HCR-FlowFISH target, and discard putative CREs which are not supported in all experimental replicates. This minimizes our Type I Error rate.

Finally we assign maximum *a posteriori* (MAP) estimates of activity to each CRE based on summits of absolute activity. First, we assume scaled activity posteriors are Gaussian and assign $\mu_{wr}$ as the midpoint of the 95% HDR of each $w$ window and $r$ replicate. Next, we linearly scale all replicates such that:

$$\max_{1 \leq w \leq W}(\mu_{w1}) = \max_{1 \leq w \leq W}(\mu_{wr}), \forall r$$

and

$$\min_{1 \leq w \leq W}(\mu_{w1}) = \min_{1 \leq w \leq W}(\mu_{wr}), \forall r.$$

To avoid expensive Bayesian inference under a model of replication we assume:

$$a^*_{wr} \mid \mathbf{l}, \mathbf{h} \sim \mathcal{N}\left(\mu^*_{wr}, \sigma^2_{wr}\right)$$

$$\mu^*_{wr} \sim \mathcal{N}\left(\zeta_w, \sigma^2\right)$$

where the asterisk indicates normalization by non-targeting controls and scaling to the first replicate. Under these assumptions, the MAP activity estimate of each bin is:

$$\mathrm{MAP}\left(\zeta_w\right) = \frac{1}{R}\sum_{r=1}^{R} \mu^*_{wr}.$$

Finally, CRE activity is determined by the most extreme MAP estimates for the $\zeta_w$'s within the CRE area.

*Reanalysis of the Prime-Flow screen for the GATA1 gene.*

We downloaded gRNA annotations and three replicates of raw CRISPRi-FlowFISH counts for the *GATA1* gene from the Open Science Framework at https://osf.io/uhnb4/ [22]. First, we remapped gRNAs to hg38 using BOWTIE1 with 0 mismatches. Next we simplified the raw counts data from a 6-way flow cytometry sort by discarding counts from the middle four expression bins; our analysis only considered counts of cells in the top and bottom 10-percentile of *GATA1* expression. Furthermore, we discarded all but the first PCR replicate from each sorting replicate. CASA was run on these simplified data using the same settings as for the analysis of HCR-FlowFISH on *GATA1*. We report putative CREs supported by at least two screen replicates.

*Functional genomic annotations and analyses*

We downloaded all functional genomic data for comparison with our screens and visualization from the ENCODE portal (https://www.encodeproject.org/) [2] with the following identifiers: ENCSR000EKS (K562 DHS), ENCSR000AKP (K562 H3K27ac), ENCSR000CIL (K562 CAGE), ENCSR000BNL (K562 SP2 ChIP-Seq). PRO-seq and GRO-seq datasets were obtained from Wang et al 2018 [54].For all analysis and visualization we used the comprehensive gencode 32 human gene annotation (https://www.gencodegenes.org/human/release_32.html) unless otherwise specified. Promoters were defined as 1000bp upstream of gencode annotated transcription start site. Transcription factor binding sites and factorbook motifs were accessed via the UCSC genome browser and the SP2 motif position weight matrix was accessed via HOCOMOCO [55,56]. K562 expression data and TPMs were calculated using the ARCH4s database [24].

*RT-qPCR*

Gene specific primers were purchased from Eton Bioscience **(Supplementary Table 2)**. We generated standard curves for each amplicon in ten-fold serial dilutions ranging in concentration from 100pM to $1\times10^{-4}$ pM using the Power SYBR Green RNA-to-CT 1-Step Kit (Thermo Fisher, 4389986) on an Applied Biosystems' QuantStudio 6 Flex. ≧3 replicates were used to determine mean Ct values. Dilutions outside the linear range were discarded.

RNA was isolated from single-guide K562 cell lines and total RNA was isolated from cells using the Qiagen's RNeasy Mini Kit (Qiagen 74106) using DTT and homogenization with a Qiashredder (79654). Subsequently, DNA was removed using the TURBO DNase Kit (Thermo Fisher, AM2238) for 30 minutes at 37°C. RNA was purified using 2X by volume Agencourt RNAClean XP SPRI beads (Beckman Coulter, A63881). 50ng of total RNA from cells was used per qPCR and target abundance determined by the previously generated standard curves.

Expression was determined using the standard curve described above to calculate fold change reduction in perturbed vs unperturbed cells. We normalized target abundance against the copy numbers of a housekeeping gene, *TBP*, across ≧3 replicates.

*HCR-FlowFISH Cellular Imaging*

Cell aliquots were taken from HCR-FlowFISH prepared samples prior to cytometric analysis and placed on a standard slide. Cells were imaged with a WideField EpiFluorescence microscope from ASI Imaging at 20x magnification with a 0.5 NA Nikon Objective. Acquisition settings were kept constant between images. A custom image analysis pipeline was created using Cell Profiler (Broad Institute) to identify cells and obtain red:green intensity quantifications per cell.

*MPRA design, experiment and analysis*

The MPRA library for the *FADS* locus was constructed as previously described [11]. Briefly, 200 bp sequences were designed that tile across hg19 at chr11:61555001-61665622 moving over the region with a 5 bp sliding window, both the forward and reverse orientation were selected for testing for a total of 44,330 sequence. All 3,108 single nucleotide polymorphisms and small indels in the 1000 genome phase 3 were also tested by centering the allele in the middle of 200 bp of flanking sequence and taking both orientations. In addition, we included 265 positive and negative control sequences selected based on their activity in previous MPRA assays. Oligos were synthesized (Agilent Technologies) as 230 bp sequences containing the 200 bp of genomic sequence and 15 bp of adaptor sequence on either end. Unique 20 bp barcodes were added by PCR along with additional constant sequence for subsequent incorporation into a backbone vector by Gibson assembly (primers MPRA_v3_F & MPRA_v3_20I_R). The oligo library was expanded by electroporation into E. coli, split immediately into 10 independent cultures and grown for 6 hours. An appropriate number of expanded cultures were selected to achieve an average of 200 CFU per oligo sequence tested. The resulting purified ΔGFP plasmid library was sequenced by Illumina 2 X 150 bp chemistry to acquire oligo-barcode pairings. The library underwent AsiSI restriction digestion, and a GFP amplicon with a minimal TATA promoter was inserted by Gibson assembly resulting in the 200 bp oligo sequence positioned directly upstream of the promoter and the 20 bp barcode falling in the 3' UTR of GFP. After expansion within E. coli the final MPRA plasmid library was sequenced by Illumina 1 X 31 bp chemistry to acquire a baseline representation of each oligo-barcode pair within the library.

100 million K562 cells were transfected with a Thermo Fisher Neon Transfection System (3 pulses, 1450V, 10 ms) using 10 million cells and 5 ug of plasmid per transfection in RPMI. Forty eight hours after transfection, cells were collected by centrifugation and washed three times with PBS prior to freezing at -80°C. RNA was extracted from frozen cell pellets using the Qiagen RNeasy Midi kit. Following DNase treatment, a mixture of 3 GFP-specific biotinylated oligos (GFP_BiotinCapture_1-3) were used to immunoprecipitated GFP transcripts using Streptavidin C1 Dynabeads (Life Technologies). Following another round of DNase treatment, cDNA was synthesized from GFP mRNA using SuperScript III and purified with AMPure XP beads. Quantitative PCR using primers specific for GFP was used to determine the cycle at which linear amplification begins for each replicate.

Replicates were diluted to approximately the same concentration based on the qPCR results, and a 14 cycle PCR with NEBNext Ultra II Q5 Master Mix was used to amplify the cDNA (primers MPRA_v3_Illumina_GFP_F & Ilmn_P5_PCR). A second round of PCR (6 cycles) was used to add Illumina sequencing adaptors and indices to the DNA/RNA replicates. The resulting MPRA barcode libraries were spiked with 5% PhiX and sequenced using Illumina single-end chemistry (with 8 bp index read) on a NextSeq 500.

Data from the MPRA was analyzed as previously described [11]. Briefly, the sum of the barcode counts for each oligo were provided to DESeq2 and replicates were median normalized followed by an additional normalization of the RNA samples to center the RNA/DNA activity distribution over a log2 fold change of zero [57]. Oligos showing differential expression relative to the plasmid input were identified by modeling a negative binomial distribution with DESeq2 and applying a false discovery rate (FDR) threshold of 1%. For sequences that displayed significant MPRA activity, a paired t-test was applied on the log-transformed RNA/plasmid ratios for each experimental replicate to test whether the reference and alternate allele had similar activity. An FDR threshold of 10% was used to identify SNPs with a significant skew in MPRA activity between alleles (allelic skew).

*Cutting library design, construction, sequencing, and analysis*
We used the 909 guides within the ~8 kb region (chr11:61635743-61643818, hg38) surrounding the *FADS1/FADS3* intergenic CRE to design a targeted subpool for use in a CRISPR cutting experiment. Oligos were synthesized by Twist Biosciences, and were amplified identically to the larger libraries. Guides were cloned into a vector bearing the catalytically active CRISPR protein (lentiCRISPR v2 Addgene: 52961). Guides were transduced and puromycin selected identically to the larger libraries. HCR was performed on cells after 10 days to allow deletions to be generated. Cells were sorted into the same 10% high and low bins as performed for the CRISPRi screens.

HCR-labeled and sorted cells were de-crosslinked and DNA isolated following the same protocol used for Illumina sgRNA libraries. The guide targeted region was amplified as a single 8.9 kb amplicon in a 50 uL reaction containing 24 uL of extracted DNA, 1 uL PrimeStar GXL polymerase, 1 uL of 10 mM dNTP, 1 uL of each primer 10 uM and 10 uL of 5X GXL buffer. Each sample was amplified with a unique primer pair containing a 5 bp barcode on the 5' end used for demultiplexing post-sequencing. Samples were equal molar pooled prior to standard SMRTBell ligation library preparation. HiFi circular consensus sequencing was performed for 30 hours on a Sequel II by the JAX Genome Technologies Laboratory group. Downstream processing was performed with the Pacbio toolset using the lima package for demultiplexing individual samples (flags: --single-side, --ccs, --window-size 5, --min-length 500) and read mapping using the minimap2 wrapper pbmm2 with standard settings for CCS reads to build 38 of the human genome [58].

Mapped reads were quantified with respect to the number of unique deletions detected at each nucleotide by selecting reads mapping to the FADS locus and supported by 2 or more CCS passes of sequencing. We next designated the genomic area within the amplicon and outside of our previously defined CREs as the control area, such that deletions in this space help define the null effect distribution similar to non-targeting gRNAs in CRISPRi screens. We then run CASA on the CRISPR cutting data using 50nt windows without extending the effects of deletions outside of the nucleotides they explicitly remove (in contrast to the effects of gRNAs in CRISPRi screen which we assume generate a wide area of effect). An odds ratio threshold of 2.0 is used to define the region of significant activity.

*Population genetic analysis, eQTLs, and UK Biobank fine-mapping*

LD ($R^2$) was reported from analysis in the 1000 Genomes projects, using individuals from Europe [59]. eQTLs were obtained from all tissues from the gTEX portal (v8.0) on 4/30/18. We used statistical fine-mapping results of 96 complex traits and diseases in the UK Biobank that we previously conducted (https://www.finucanelab.org/data) [45]. Briefly, we conducted GWAS and statistical fine-mapping in the UK Biobank using up to 361,194 white British individuals. We computed association statistics for the variants with INFO > 0.8, MAF > 0.01% (except for rare coding variants with MAC > 0), and HWE p-value > 1e-10 using SAIGE (for case-control studies) [60] or BOLT-LMM (for continuous traits) [61] with the covariates including top 20 PCs, sex, age, $age^2$, sex * age, sex * $age^2$, and dilution factor where appropriate. Statistical fine-mapping was performed using FINEMAP v1.3.1 [62,63] and susieR v0.8.1.0521 [64] with the maximum number of causal variants specified as 10. We defined a region based on a 3 Mb window around a lead variant and then merged any regions that overlapped. We used summary statistics from the GWAS and in-sample LD matrices which were calculated from imputed dosages for individuals included in each GWAS using LDstore v2.0b.

*Guide design optimization analysis*

We subsetted the number of guides used to analyze CRE interactions with *FADS1* and determined the impact of guide density on peak calling with CASA. We subsetted guides by applying a 100nt sliding window to the *FADS* locus and selecting a specified fraction of guides targeting each window for removal, deterministically. Guides with lower predicted cutting specificity scores were removed first.

*Data availability*

All raw CRISPRi and CRISPR cutting screening data and MPRA data as well as process files have been uploaded to the ENCODE Portal and is currently being processed for release via the portal and in a linked GEO accession.

Software used to run CASA and generate scores is available at https://github.com/sjgosai/casa. Track hubs are available for each locus screened at the following links:

https://genome.ucsc.edu/s/skr2/GATA_HCR
https://genome.ucsc.edu/s/skr2/CD164_HCR
https://genome.ucsc.edu/s/skr2/ERP29_HCR
https://genome.ucsc.edu/s/skr2/LMO2_HCR
https://genome.ucsc.edu/s/skr2/NMU_HCR
https://genome.ucsc.edu/s/skr2/MEF2C_HCR
https://genome.ucsc.edu/s/skr2/FADS_HCR

***Competing Interests***

PCS is a co-founder of and consultant to Sherlock Biosciences and Board Member of Danaher Corporation.

***References***

44. Bycroft, C. *et al.* The UK Biobank resource with deep phenotyping and genomic data. *Nature* **562**, 203–209 (2018).
45. Ulirsch, JC. & Kanai, M., et al. A compendium of candidate causal variants for 96 complex traits and gene expression in 49 tissues. *(In Preperation)*
46. Doench, J. G. *et al.* Optimized sgRNA design to maximize activity and minimize off-target effects of CRISPR-Cas9. *Nat. Biotechnol.* **34**, 184–191 (2016).
47. Langmead, B., Trapnell, C., Pop, M. & Salzberg, S. L. Ultrafast and memory-efficient alignment of short DNA sequences to the human genome. *Genome Biol.* **10**, R25 (2009).
48. Hsu, P. D. *et al.* DNA targeting specificity of RNA-guided Cas9 nucleases. *Nat. Biotechnol.* **31**, 827–832 (2013).
49. Morgens, D. W. *et al.* Genome-scale measurement of off-target activity using Cas9 toxicity in high-throughput screens. *Nat. Commun.* **8**, 15178 (2017).
50. Wang, T., Lander, E. S. & Sabatini, D. M. Viral Packaging and Cell Culture for CRISPR-Based Screens. *Cold Spring Harb. Protoc.* **2016**, db.prot090811 (2016).
51. Kruschke, J. K. Rejecting or Accepting Parameter Values in Bayesian Estimation. *Advances in Methods and Practices in Psychological Science* **1**, 270–280 (2018).
52. Salvatier, J., Wiecki, T. V. & Fonnesbeck, C. Probabilistic programming in Python using PyMC3. *PeerJ Comput. Sci.* **2**, e55 (2016).
53. Hoffman, M. D. & Gelman, A. The No-U-Turn Sampler: Adaptively Setting Path Lengths in Hamiltonian Monte Carlo. *J. Mach. Learn. Res.* **15**, 1593–1623 (2014).
54. Wang, J. *et al.* Nascent RNA sequencing analysis provides insights into enhancer-mediated gene regulation. *BMC Genomics* **19**, 633 (2018).
55. Wang, J. *et al.* Factorbook.org: a Wiki-based database for transcription factor-binding data generated by the ENCODE consortium. *Nucleic Acids Res.* **41**, D171–6 (2013).
56. Kulakovskiy, I. V. *et al.* HOCOMOCO: towards a complete collection of transcription factor binding models for human and mouse via large-scale ChIP-Seq analysis. *Nucleic Acids Res.* **46**, D252–D259 (2018).
57. Love, M. I., Huber, W. & Anders, S. Moderated estimation of fold change and dispersion for RNA-seq data with DESeq2. *Genome Biol.* **15**, 550 (2014).
58. Li, H. Minimap2: pairwise alignment for nucleotide sequences. *Bioinformatics* **34**, 3094–3100 (2018).
59. Sudmant, P. H. *et al.* An integrated map of structural variation in 2,504 human genomes. *Nature* **526**, 75–81 (2015).
60. Zhou, W. *et al.* Efficiently controlling for case-control imbalance and sample relatedness in large-scale genetic association studies. *Nat. Genet.* **50**, 1335–1341 (2018).
61. Loh, P.-R., Kichaev, G., Gazal, S., Schoech, A. P. & Price, A. L. Mixed-model association for biobank-scale datasets. *Nat. Genet.* **50**, 906–908 (2018).
62. Benner, C. *et al.* FINEMAP: Efficient variable selection using summary data from genome-wide association studies. *Bioinformatics* **32**, 1493–1501 (2016).
63. Benner, C., Havulinna, A. S., Salomaa, V., Ripatti, S. & Pirinen, M. Refining fine-mapping: effect sizes and regional heritability. *bioRxiv* 318618 (2018) doi:10.1101/318618.
64. Wang, G., Sarkar, A., Carbonetto, P. & Stephens, M. A simple new approach to variable selection in regression, with application to genetic fine-mapping. *bioRxiv* 501114 (2019) doi:10.1101/501114.

***Supplementary Tables***

**Supplementary Table 1:** HCR probe details and targets.
**Supplementary Table 2:** Sequencing primers, qpcr primers, MPRA primers, guides.
**Supplementary Table 3:** CRISPRi HCR-FlowFISH screen results.
**Supplementary Table 4:** Cutting HCR-FlowFISH screen results.
**Supplementary Table 5:** CASA CRE calls and activity scores
**Supplementary Table 6:** MPRA results.
**Supplementary Table 7:** FADS locus SNPs, GWAS, and fine-mapping results.

***Supplementary Figures***

**Supplemental Fig. 1** Comparison of transcript abundance detection between HCR-FlowFISH and PrimeFlow.

**Supplemental Fig. 2** CRISPRi induction, sorting schema, and construction of CASA (CRISPR Activity Screen Analysis) a generative model of CRE activity.

**Supplemental Fig. 3** HCR-FlowFISH screens display high similarity and increased sensitivity compared to growth screens at the *GATA1* locus.

**Supplemental Fig. 4** HCR-FlowFISH and CASA enhance selectivity of CRISPRi screens at the GATA1 locus.

**Supplemental Fig. 5.** Permissive CASA analysis identifies a CRE with modest activity on NMU and links a GWAS regulatory variant to MEF2C.

**Supplemental Fig. 6.** HCR-FlowFISH produces reproducible effects in CRISPRi screens at the FADS locus.

**Supplemental Fig. 7.** Dense gRNA-tiling at the FADS locus informs guide design and CRE calling limitations.

**Supplemental Fig. 8.** HCR-FlowFISH and CASA reveal complex CRE sharing at the FADS locus.

**Supplemental Fig. 9.** CREs identified by CASA and HCR-FlowFISH show signatures of cis-regulatory elements.

**Supplemental Fig. 10.** CRE activity is weakly correlated with DHS intensity and not well correlated with distance to target gene**.**

**Supplemental Fig. 11.** HCR-FlowFISH identifies elements where variants have opposing regulatory effects on different genes.

**Supplemental Fig. 12.** Functional characterization nominates rs174466 as *FADS3* CRE-activity altering SNP
